# Viral delivery of an RNA-guided genome editor for transgene-free germline editing in *Arabidopsis*

**DOI:** 10.1101/2024.07.17.603964

**Authors:** Trevor Weiss, Maris Kamalu, Honglue Shi, Zheng Li, Jasmine Amerasekera, Zhenhui Zhong, Benjamin A. Adler, Michelle Song, Kamakshi Vohra, Gabriel Wirnowski, Sidharth Chitkara, Charlie Ambrose, Noah Steinmetz, Ananya Sridharan, Diego Sahagun, Jillian F. Banfield, Jennifer A. Doudna, Steven. E. Jacobsen

**Affiliations:** Department of Molecular, Cell and Developmental Biology, University of California at Los Angeles, Los Angeles, CA 90095, USA; Innovative Genomics Institute, University of California, Berkeley, CA 94720 USA; Department of Bioengineering, University of California, Berkeley, CA 94720 USA; Department of Molecular and Cell Biology, University of California, Berkeley, CA 94720, USA; Department of Biotechnology, Sichuan University, Chengdu, China; Howard Hughes Medical Institute, University of California, Berkeley, CA 94720, USA; Howard Hughes Medical Institute (HHMI), University of California at Los Angeles, Los Angeles, CA 90095, USA; Department of Earth and Planetary Science, University of California, Berkeley, CA 94720, USA; Department of Environmental Science, Policy and Management, University of California, Berkeley, CA 94720, USA; University of Melbourne, Melbourne, Australia; California Institute for Quantitative Biosciences (QB3), University of California, Berkeley, CA 94720, USA; Department of Chemistry, University of California, Berkeley, CA 94720, USA; MBIB Division, Lawrence Berkeley National Laboratory, Berkeley, CA 94720, USA; Gladstone Institutes, University of California, San Francisco, CA 94158, USA

## Abstract

Genome editing is transforming plant biology by enabling precise DNA modifications. However, delivery of editing systems into plants remains challenging, often requiring slow, genotype-specific methods such as tissue culture or transformation. Plant viruses, which naturally infect and spread to most tissues, present a promising delivery system for editing reagents. But most viruses have limited cargo capacities, restricting their ability to carry large CRISPR-Cas systems. Here, we engineered tobacco rattle virus (TRV) to carry the compact RNA-guided TnpB enzyme ISYmu1 and its guide RNA. This innovation allowed transgene-free editing of *Arabidopsis thaliana* in a single step, with edits inherited in the subsequent generation. By overcoming traditional reagent delivery barriers, this approach offers a novel platform for genome editing, which can greatly accelerate plant biotechnology and basic research.

---

Programmable RNA-guided endonucleases including CRISPR-Cas9 are driving advances in genome editing for both fundamental research and biotechnology. The ability to genetically modify plant genomes has allowed for the creation of rationally designed phenotypes. However, efficient delivery of genome editing reagents to plants remains a major challenge. The most common strategy is to encode RNA-guided genome editors (e.g. CRISPR-Cas enzymes) within transgenes and use tissue culture and plant transformation approaches to make transgenic plants, after which genetic crosses are required to remove the transgenic material but retain the edits ^1–3^. However, current plant transformation methods are limited to specific plant species and genotypes, often require considerable time, resources and technical expertise, and can cause unintended changes to the genome and epigenome ^1^.

One approach to circumvent these limitations is to use plant viral vectors to deliver genome editing reagents. For example, several viral vectors have been engineered to encode guide RNAs (gRNAs) for delivery to transgenic plants already expressing Cas9, resulting in somatic and germline editing and transmission of edits to the next generation ^4–7^. Because plants have evolved mechanisms to restrict viral infection of meristems and germ cells, most viruses are rarely sexually transmitted ^8^. However, transient invasion of meristem cells by viral RNAs encoding gRNAs can allow these cells to be edited, and for these edits to be seed transmissible ^4–7^. While these approaches represent significant advances, they still require the use of transgenic plants to express the nuclease protein.

A strategy to avoid the need for transgenic plant materials has been the use of viral vectors with large cargo capacities, capable of expressing entire editing systems (e.g. Cas9 and the gRNA). This approach has been met with some success, however it still requires plant regeneration steps because these viruses do not cause germline editing and heritability of the edits ^9–12^. On the other hand, encoding entire CRISPR systems in viruses that are capable of germline transmission has not been possible because of their limited cargo capacity ^4–7,13^.

To overcome this cargo size limit, we explored the potential of TnpB, a class of ultracompact RNA-guided endonucleases (∼400 amino acids) ^14–16^, to be encoded in a plant RNA viral vector. As ancestors of Cas enzymes, TnpBs similarly utilize a programmable RNA guide, called an omega RNA (ωRNA), to be directed to any target site and induce genome edits. Previously, TnpBs ISDra2, ISYmu1 and ISAam1 were shown to be capable of targeted genome editing in mammalian cells, and ISDra2 and ISYmu1 in monocot rice plant cells ^14,17–19^. Here, we tested the ISDra2, ISYmu1 and ISAam1 TnpBs for genome editing in the dicot plant, *Arabidopsis*. Given the single cargo site in the TRV vector that is typically used, we sought to express both the TnpB protein and its guide RNA within the same mRNA transcript under a single promoter, similar to their natural expression arrangement ^14–16^.

To test the activities of TnpB and its gRNA encoded in a single transcript, we first expressed these three TnpBs and assessed their RNA-guided plasmid interference activities in bacteria. We co-expressed the TnpB and gRNA from the same promoter as a single transcript, maintaining their natural sequences without codon optimization. We compared two configurations of the 3’-guide region: one extended continuously without a terminator to mimic the natural TnpB condition, and another capped by the hepatitis delta virus (HDV) ribozyme, as previously used in bacteria ^14^ (Extended Data Fig. 1). Our results showed that without the HDV ribozyme, only ISDra2 demonstrated plasmid interference activity whereas with the HDV ribozyme, all three TnpBs exhibited robust activity at both 26°C and 37°C (Fig. 1a, Extended Data Fig. 2). These findings revealed that single transcript expression cassettes with an HDV ribozyme sequence at the 3’ end are capable of cleaving plasmid DNA in bacteria.

**Figure 1:**
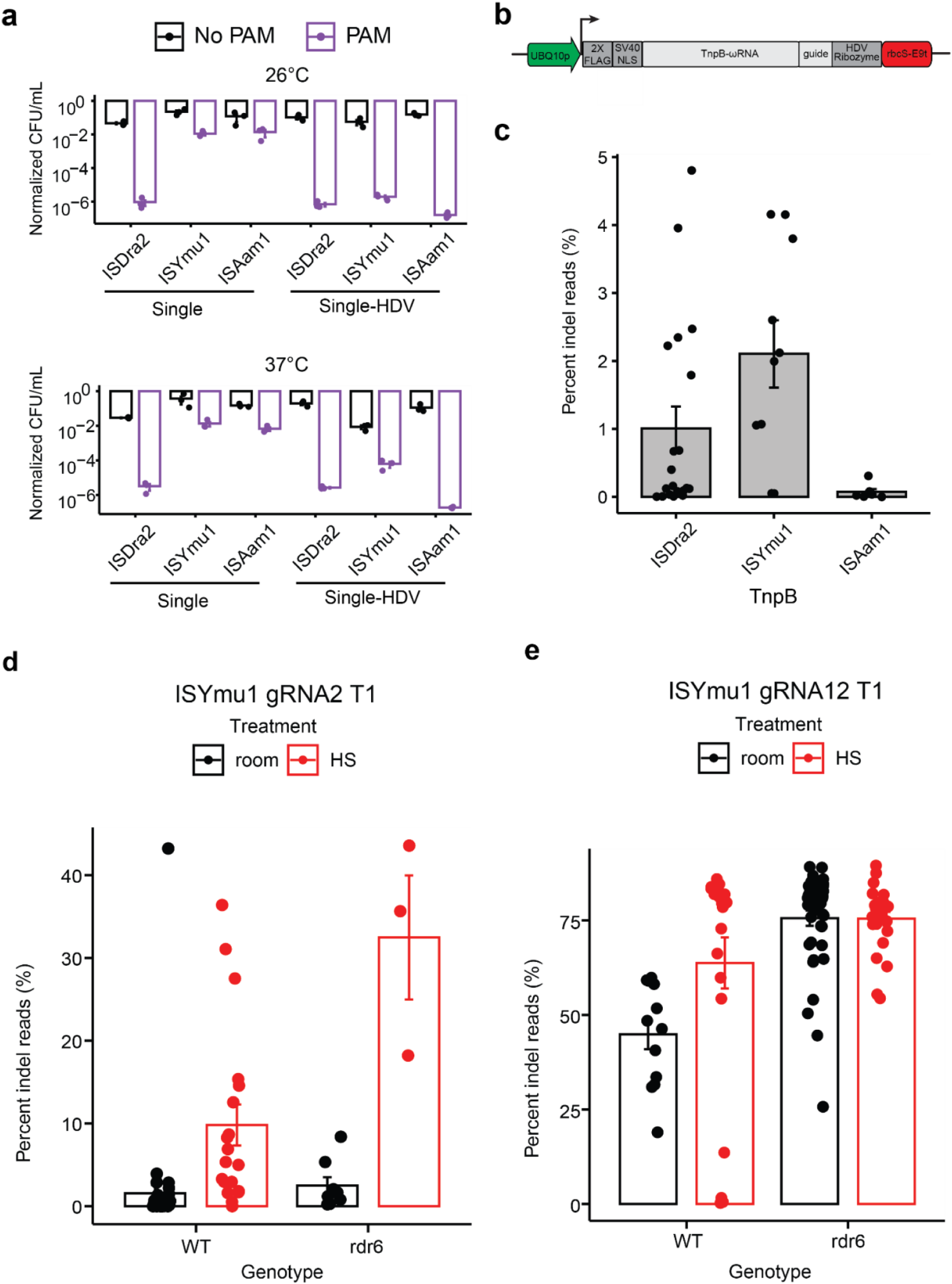
Expression of TnpB and guide RNA in a single transcript for plant genome editing. **(a)** Barplots of interference assay testing the single transcript expression TnpB vectors for cleavage in E. coli. The barplot on the top displays data from experiments performed at 26**°**C and the barplot below depicts data from experiments performed at 37**°**C. Black bars indicate absence of a PAM on the target plasmid and purple bars indicate the presence of a PAM on the target plasmid. The Y-axis is a log10 scale of the normalized colony forming units (CFU)/milliliter (mL). The X-axis displays the three TnpBs tested using the single expression transcript design without a HDV ribozyme or the single expression design with a HDV ribozyme. The standard error of the mean (SEM) was calculated for each experiment. **(b)** Schematic of the single expression transcript TnpB-ωRNA plasmid design used for plant genome editing. The green arrow symbolizes the At*UBQ10* promoter; the dark gray boxes indicate the 2X-FLAG, SV40 NLS and HDV ribozyme sequences; the light gray boxes indicate the TnpB-ωRNA and guide sequences; the red box symbolizes the rbcS-E9 terminator; the black arrow indicate the orientation of the TnpB-ωRNA expression cassette. **(c)** Barplot displaying the average editing frequency for protoplast experiments using ISDra2, ISYmu1 and ISAam1. Each dot represents the average editing efficiency of a gRNA from Extended Data Fig. 3a. The Y-axis indicates the editing efficiencies (percent indel reads (%)). The X-axis displays the ISDra2, ISYmu1 and ISAam1 TnpBs tested. The standard error of the mean (SEM) was calculated for each experiment. **(d-e)** ISYmu1 somatic editing in T1 transgenic plants. Panels d and e display a barplot for ISYmu1 gRNA2 and ISYmu1 gRNA12, respectively. Each dot indicates a single T1 transgenic plant. The room and HS treatments stand for room temperature and heat shock plant growth conditions, respectively. The genotypes are plotted along the X-axis and the editing efficiencies (percent indel reads (%)) are plotted on the Y-axis. The standard error of the mean (SEM) was calculated for each experiment. WT is an abbreviation for wild type, and *rdr6* is an abbreviation for a mutation in the *RNA-DEPENDENT RNA POLYMERASE 6* gene.

To test the single expression cassette for targeted genome editing in *Arabidopsis*, we used the *AtUBQ10* promoter to drive expression of the TnpB-ωRNA and a gRNA targeting the *PHYTOENE DESATURASE3* (*AtPDS3*) gene region, followed by the HDV ribozyme and rbcS-E9 terminator (Fig. 1b). We tested twenty ISDra2 sites, ten ISYmu1 sites, and seven ISAam1 sites for editing capabilities in *Arabidopsis* protoplast cells (Supplementary Table 1) ^20^. ISDra2 and ISYmu1 demonstrated active editing ranging from 0-4.8% and 0.1-4.2%, respectively, as measured by next generation amplicon sequencing (amp-seq) (Extended Data Fig. 3a). ISAam1 was much less active, ranging from 0-0.3% editing efficiency (Extended Data Fig. 3a). On average, we observed editing efficiencies of 1% for ISDra2, 2.1% for ISYmu1 and 0.1% for ISAam1 (Fig. 1c). In line with previous reports, the DNA repair profiles consisted of deletion-dominant repair outcomes for all three TnpBs (Extended Data Fig. 3b) ^14,17,18^. These data demonstrate that ISDra2, ISYmu1 and ISAam1 are all capable of targeted genome editing in *Arabidopsis* plant cells using the single transcript expression design.

To evaluate TnpB-mediated editing in transgenic plants we selected ISYmu1, as it demonstrated the highest average editing efficiency in *Arabidopsis* protoplast cells and was shown to exhibit no off-target editing in rice ^18^. Two gRNAs with the most active editing were selected, each targeting a unique genomic context. gRNA2 targeted the coding region of *AtPDS3* whereas gRNA12 targeted the promoter region directly upstream of the *AtPDS3* gene. Transgenic plants were created via standard floral dip transformation utilizing the same plasmids as for the protoplast experiments ^21^. To test for sensitivity to temperature, transgenic plants expressing ISYmu1 were either grown at room temperature or subjected to a heat shock treatment. We tested editing in wild type (WT) plants, as well as in the *rna dependent rna polymerase 6* (*rdr6*) mutant which is known to reduce transgene silencing ^22^. Analysis using amp-seq revealed an average editing efficiency of 1.6% and 2.5% for gRNA2 in WT and *rdr6*, respectively (Fig. 1d). Analysis of gRNA12 revealed greater editing than gRNA2, averaging 44.9% editing in WT and 75.5% in *rdr6* (Fig. 1e). Comparison of editing efficiency in the plants grown at room temperature with those that received the heat shock treatment revealed a preference for increased temperature for both target sites in the WT background, demonstrating 6.3-fold and 1.4-fold increases in editing for gRNA2 and gRNA12, respectively (Fig. 1d, e). In *rdr6* we observed a 13-fold increase in editing for gRNA2, but little change in editing for gRNA12 (Fig. 1d, e). These data demonstrate that ISYmu1, encoded as a transgene, is capable of performing efficient genome editing in *Arabidopsis* plants, and that heat treatment and the *rdr6* silencing mutant can be used to increase editing efficiency.

Encouraged by the ISYmu1 activity in transgenic *Arabidopsis* plants, we next tested ISYmu1 for TRV-mediated genome editing. TRV is a bipartite RNA virus composed of TRV1 and TRV2 (Fig. 2a). Previous work has shown that the TRV2 RNA can be engineered by inserting a cargo expression cassette downstream of the pea early browning virus promoter (pPEBV) (Fig. 2a) ^23,24^. To test ISYmu1 for genome editing capabilities via TRV delivery to *Arabidopsis,* we engineered two TRV2 Cargo Architectures. In TRV2 Architecture_A, the tRNA^Ileu^ was directly downstream of the TnpB and gRNA sequences (Fig. 2a). In TRV Architecture_B, we included an HDV ribozyme sequence between the guide and tRNA^Ileu^ sequences.(Fig. 2a). We included tRNA^Ileu^ in both designs as it was previously shown to promote systemic TRV movement and transmission of edited alleles to the next generation ^23,24^.

**Figure 2:**
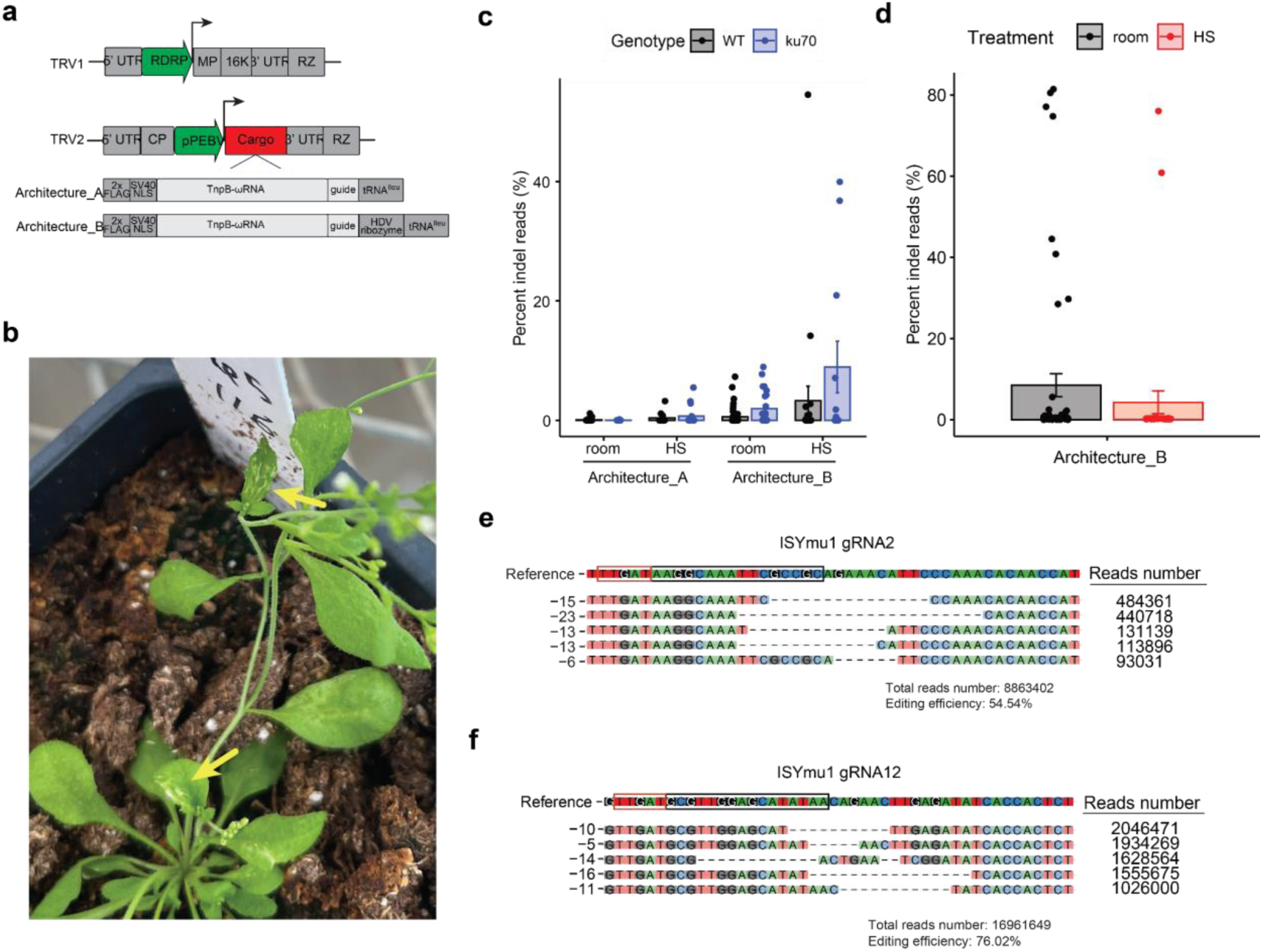
Somatic editing of *Arabidopsis* using TRV to deliver ISYmu1 TnpB and guide RNA. **(a)** Schematic of the TRV1 and TRV2 plasmids. Green arrows indicate the RNA dependent RNA polymerase (RDRP) and pea early browning virus (pPEBV) promoters for TRV1 and TRV2, respectively; the gray boxes in TRV1 and TRV2 indicate the native TRV components; the red Cargo box in TRV2 indicates the location of either Architecture_A or Architecture_B; below TRV2 are schematics of the components, Architecture_A or Architecture_B, cloned into the TRV2 Cargo slot. **(b)** Representative picture of a plant displaying white sectors about three weeks after TRV delivery. The yellow arrows indicate leaves containing white sectors. **(c)** Barplot displaying the somatic editing efficiencies for ISYmu1 gRNA2 in wild type (WT) and *ku70* genetic backgrounds. WT samples are black and *ku70* samples are blue. The TRV2 Cargo Architectures are plotted along the X-axis with either room or HS treatment. The room and HS along the X-axis stand for room temperature and heat shock plant growth conditions, respectively. The Y-axis indicates the editing efficiencies (percent indel reads (%)). Each dot represents an individual plant that underwent agroflood TRV delivery. The standard error of the mean (SEM) was calculated for each experiment. **(d)** Barplot displaying the somatic editing efficiencies for ISYmu1 gRNA12 in wild type (WT). Room temperature treatment (room) samples are black and heat shock (HS) samples are red. The TRV2 Cargo Architecture_B is plotted along the X-axis. The Y-axis indicates the editing efficiencies (percent indel reads (%)). Each dot represents an individual plant that underwent agroflood TRV delivery. The standard error of the mean (SEM) was calculated for each experiment. **(e-f)** Panel e displays the DNA indel repair profile for an individual WT plant that underwent delivery of TRV Cargo Architecture_B with ISYmu1 gRNA2 under the heat shock treatment. Panel f shows the DNA indel repair profile for an individual WT plant that underwent delivery of TRV Cargo Architecture_B with ISYmu1 gRNA12 under the heat shock treatment. The top five most common indel types are listed on the left. The read counts for each indel are listed on the right. The PAM is identified by the red box, and the target site is outlined by the black box, in the Reference sequence. The total read number and editing efficiency are listed below each indel profile.

First, we evaluated TRV-mediated editing potential with gRNA2 using both TRV2 Architecture_A and Architecture_B. gRNA2 was selected because it targets the *AtPDS3* coding sequence, enabling easy phenotypic screening for editing due to white photobleaching of cells containing biallelic mutations ^23,24^. We delivered TRV vectors to both WT and the *ku70* genetic mutant. *Ku70* plays a role in the nonhomologous end joining (NHEJ) double strand break (DSB) repair pathway ^25^. ISYmu1-mediated editing efficiency should be greater in the *ku70* genotype if double stranded breaks generated by ISYmu1 are repaired through NHEJ. Each TRV2 plasmid was co-delivered with the TRV1 plasmid to *Arabidopsis* plants using the agroflood method ^24^. White speckles were observed on some of the leaves around three weeks post agroflooding, suggesting that sectors of cells contained biallelic mutations in the target *AtPDS3* gene (Fig. 2b). Amp-seq analysis revealed an average of 0.1% and 0% editing efficiency in leaf tissue of WT and *ku70* plants agroflooded with TRV2 Architecture_A and grown under room temperature, respectively (Fig. 2c). For the heat shock treated plants, we observed an average editing efficiency of 0.4% in WT and 0.7% in *ku70* plants agroflooded with TRV2 Architecture_A (Fig. 2c). Using TRV2 Architecture_B we observed an average 0.6% and 2% editing in WT and *ku70*, respectively, for the room temperature grown plants. For the plants that received TRV Architecture_B and a heat shock, we observed an average editing efficiency of 3.3% in WT and 8.9% in *ku70* (Fig. 2c). These results show that Architecture_B, containing the HDV ribozyme, generated higher editing than Architecture_A, and that the *ku70* mutant can enhance editing efficiency.

Next, we tested gRNA12 utilizing the TRV2 Architecture_B, since this architecture demonstrated the highest levels of editing for gRNA2. Using the same agroflood TRV delivery method, we observed an average 8.51% and 4.27% editing efficiency in room temperature and heat treatment growth conditions, respectively (Fig. 2d). Further, 6/57 plants displayed editing greater than 40%, with four greater than 75%, when the room temperature treatment was used (Fig. 2d). Again, analysis of the repair outcomes showed deletion dominant profiles for ISYmu1 gRNA2 and gRNA12 (Fig. 2e, f).

To test for transmission of edited alleles to the next generation, we first screened the progeny of a WT plant showing 54.54% somatic editing using the TRV2 Architecture_B design with gRNA2 that underwent heat shock treatment. In total, 2318 seeds were sown on ½ MS plates containing 3% sucrose. After ten days, 68 albino seedlings were observed, suggesting biallelic mutations in the *PDS3* gene (Fig. 3a). To confirm *AtPDS3* was mutated we performed Sanger sequencing on the two white seedlings shown in Fig. 3a, which revealed both plants to be homozygous for a 4bp frame-shift deletion (Fig. 3b). To further characterize transmission of edited alleles, amp-seq on 209 seedlings (41 albino and 168 green) showed that all of the albino seedlings contained biallelic mutations, with the majority of mutations being the 4bp deletion observed in Fig. 3b (Supplementary Table 2). Of the 168 green seedlings, eight were heterozygous (4bp deletion/WT) (Supplementary Table 2).

**Figure 3:**
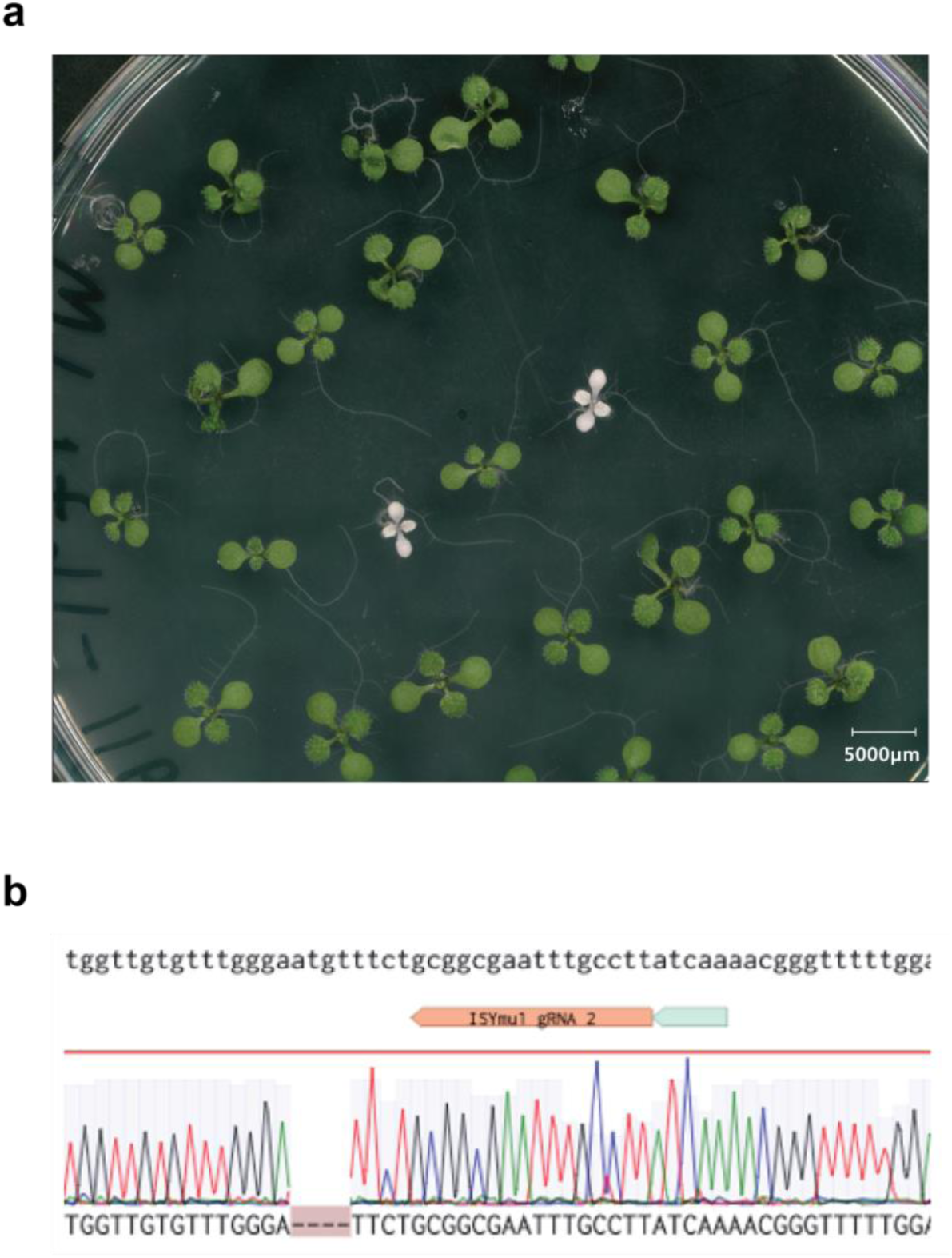
Inheritance of edited alleles. **(a)** Representative image of albino and green progeny seedlings from a WT plant showing 54.54% somatic editing using the TRV2 Architecture_B design with gRNA2 that underwent heat shock treatment. **(b)** Sanger sequencing trace file screenshot from one of the albino plants in Fig 3a. The sequence at the top is the wild type reverse complement; below that are the ISYmu1 gRNA2 target and PAM (gray box); the ab1 trace file displays a homozygous 4bp deletion.

Next, we characterized transmission of edited alleles from two individual lines, plant 54 (80.5% somatic editing) and plant 69 (77.1% somatic editing), that underwent agroflood using gRNA12 TRV2 Architecture_B with the room temperature condition. Using Sanger sequencing, we analyzed the genotypes of 148 and 75 progeny seedlings from plants 54 and 69, respectively. Sanger sequencing analysis of the progeny from plant 54 revealed 27 (18%) biallelic and 25 (17%) monoallelic edited plants (Supplementary Table 2, Extended Data Fig. 4a)^26^. For plant 69 we observed higher transmission of edited alleles, totaling 32 (43%) biallelic and 15 (20%) monoallelic edited plants (Supplementary Table 2, Extended Data Fig. 4b) ^26^. These data demonstrate the heritability of edits generated via TRV-delivery of ISYmu1 at two distinct target sites.

It has been demonstrated that TRV is not transmitted to the next generation following agroflood inoculation of plants ^23,24^. To confirm the TRV was not present in the progeny of a TRV-infected plant, RT-PCR was performed on five albino plants harboring homozygous 4bp deletions at *AtPDS3*. Consistent with the literature, TRV was not detected in any of the albino plants (Extended Data Fig. 5) ^23,24^. These data indicate that TRV-mediated biallelic edits using ISYmu1 are heritable and virus-free.

To evaluate off-target editing we surveyed three individual albino plants harboring biallelic mutations generated by ISYmu1 TRV2 Architecture_B gRNA2. Whole genome sequencing was performed to generate an average 770x coverage, with greater than 99% of the genome covered by mapped reads (Supplementary Table 3). In all three samples, we confirmed the targeted mutations in the *AtPDS3* gene, as previously identified using amp-seq. Additionally, we found a large number of variant differences compared to the Col-0 reference genome both in the control and the edited plants (Supplementary Table 4), suggesting that most of the variants detected are due to spontaneous mutations present in our lab strain of *Arabidopsis*. To screen for variants potentially caused by ISYmu1 off-target editing, all variants in the edited plants were filtered with variants already present in the control background. Variants with coverage lower than 30-fold were also filtered out. The remaining variants were checked manually for any false positive variant calling. In the three albino plants we sequenced, only 5, 5, and 4 variants were detected, and these variants are all outside the predicted potential off-target sites based on sequence similarity to the PDS3 gRNA2 sequence (Supplementary Tables 4, 5) ^27^. In line with ISYmu1 off-target analysis reported in rice and human cells ^17,18^, these data further demonstrate the high target site specificity of ISYmu1.

A long-term goal of plant scientists has been the development of fast and easy means of editing plant genomes without the need for tissue culture and transgenesis. Here, we developed an approach utilizing the ultracompact site-specific TnpB genome editor, ISYmu1, together with tobacco rattle virus, for heritable plant genome editing. These results should accelerate high throughput genome editing for both basic and applied research. We anticipate this approach will be applicable to other novel TnpBs, various viral vectors and a number of plant species for genome editing. Recent work has uncovered many TnpB systems from diverse microbial sources, including enzymes with unique PAM sequence specificities ^28^, which can increase the range of target DNA sequences that could be edited using this approach. The TRV virus used in this study has a broad host range of over 400 species, including many solanaceous plants such as tomato, ornamental plants, and other crops ^29^. Additionally, plant viruses with similar cargo capacities such as Potato Virus X and Barley Stripe Mosaic Virus are likely to be amenable to this approach since it has been demonstrated they are capable of viral-mediated heritable gene editing by delivering the gRNA to a Cas9-expressing transgenic plant ^30^. In addition to being an important tool for crop biotechnology, viral delivery of TnpBs could enable high throughput CRISPR screens in model plant species such as *Arabidopsis*, further unlocking their potential for genetic discovery.

## Materials and Methods

### Plasmids used in this study

Plasmids used for bacterial assay were generated as follows. The single expression cassette containing TnpB and ωRNA sequences were synthesized as geneblocks from IDT (Integrated DNA Technologies), and were golden-gate cloned using BsmbI restriction enzyme (cat E1602S) into a vector (chloramphenicol resistance) under a single tetracycline-inducible promoter (TetR/pTet) to make the TnpB-ωRNA plasmid. Target sites with various PAM sequences and target sites were golden-gate cloned with BBsI restriction enzyme(cat R3539S) into a vector (ampicillin/carbenicillin resistance).

Plasmids were generated for protoplast and floral dip experiments in a two-step cloning strategy. In step one, the ISDra2, ISYmu1, and ISAam1 protein coding sequences and their ωRNAs were synthesized as geneblocks by IDT (Integrated DNA Technologies). Then, starting with the pC1300_pUB10_pcoCASphi_E9t_MCS_version2 vector (Li et al. 2023), we used NEBuilder HiFi DNA Assembly (catE2621) and PCR to assemble the TnpB-ωRNA geneblocks into plant expression vectors with a toxic *ccdB* insert flanked by PaqCI sites immediately downstream of the ωRNA scaffold and preceding an HDV ribozyme sequence. The HiFi reactions were then transformed into One Shot *ccdB* Survival 2 T1R Competent Cells (catA10460) to obtain the pMK003 (ISDra2), pMK025 (ISYmu1), and pMK024 (ISAam1) intermediate vectors for facile guide sequence cloning (Supplementary Table 6). In step two, guide sequences were synthesized as individual top and bottom strands with 4 base pair overhangs from IDT, phosphorylated and annealed, and then used for golden-gate assembly using the NEB PaqCI (catR0745) enzyme (Supplementary Table 7). When transformed into NEB 10-beta Competent E. coli (catC3019) vectors that still contained the *ccdB* gene would kill the cells, leaving behind only the transformants possessing successfully assembled TnpB plant expression vectors harboring a guide RNA sequence. A full list of all plasmids can be found in Supplementary Table 6.

TRV vectors were created with the pDK3888 TRV2 plasmid as a base vector ^24^. NEBuilder HiFi DNA Assembly (catE2621) was used to clone the ISYmu1 gRNA2 Architecture_A, ISYmu1 gRNA2 Architecture_B, and ISYmu1 gRNA12 Architecture_B into the TRV2 cargo slot. First, pDK3888 was digested using NEB ZraI (catR0659), NEB PmlI (catR0532), and NEB Quick CIP (catM0525) overnight, and purified using Qiagen QiaQuick purification column (cat28104). Next, three PCR reactions were performed to amplify the fragments needed for NEBuilder HiFi DNA Assembly, followed by purification using Qiagen QiaQuick purification column (cat28104) (Supplementary Table 8). Then, the digested and purified pDK3888 plasmid, and purified PCR fragments were used to assemble the final TRV2 plasmid using NEBuilder HiFi DNA Assembly according to the manufacturer’s protocol. Finally, the NEBuilder HiFi DNA Assembly reaction was transformed into NEB 10-beta Competent E.coli (catC3019). Correct plasmids were confirmed using Primordium whole plasmid sequencing. Plasmids and their descriptions can be found in Supplementary Table 6.

### Bacterial interference assay

For the bacterial interference assay, we co-transfected 100 ng of the TnpB-ωRNA plasmid and 100 ng of the target plasmid to 33 µl of NEB 10-beta electrocompetent E. coli cells (cat C3020K). Specifically, the target plasmid contains a target site either flanking the canonical PAM (TTGAT for ISYmu1 and ISDra2 and TTTAA for ISAam1) or flanking a non-canonical PAM (GGGGG). The cells were recovered in 1 ml of NEB 10-Beta Stable/Outgrowth media(cat B9035S) for 1 hour. Following recovery, a series of 5-fold dilutions of the recovery culture were prepared. Each dilution (5 μl) was spot-plated onto LB-Agar plates containing double antibiotics (34 μg/ml chloramphenicol, 100 μg/ml carbenicillin, and 2 nM anhydrotetracycline) and onto control plates with a single antibiotic (34 μg/ml chloramphenicol and 2 nM anhydrotetracycline). If no colonies were visible on the serial dilution plates, 400 μl of the 1 ml recovery culture was plated entirely on the double antibiotic plate to enhance detection sensitivity. Plates were left overnight at either 26°C or 37°C, and colony-forming units (CFU) were counted on all plates the next morning. The normalized CFU were calculated by taking the ratio of CFU on the double antibiotic plates to the CFU on the single antibiotic plates. The normalized CFU in the canonical PAM conditions were compared to the one in the non-canonical PAM conditions. Experiments were performed in triplicate.

### Plant materials and growth conditions

For protoplast preparation, *Arabidopsis* Columbia ecotype (Col-0) seeds were suspended in a 0.1% agarose solution and kept at 4**°**C in the dark for three days to stratify. Following stratification, seeds were planted on Jiffy pucks and grown under a 12-h light/12-h dark photoperiod, with low light condition for 3 to 4 weeks ^20^.

For the creation of transgenic plants, the *Arabidopsis* Col-0 ecotype was used. The *ku70* (SALK_123114) genotype was provided from Feng Zhang lab at the University of Minnesota. The *rdr6* genotype was created using CRISPR-Cas9, resulting in a 616 bp deletion in the gene body of *rdr6*. Floral dip transformation was performed according to the protocol as previously outlined using the Agl0 agrobacterium strain ^21^. Transgenic T1 plants were screened using ½ MS plates with 40 μg/mL hygromycin B under a 16-/8-h light/dark cycle at 23**°**C. After one week, transgenic seedlings that passed selection were transferred to soil and moved to a greenhouse (23°C) for the rest of their life cycle.

For agroflood experiments, sterilized seeds were sown on ½ MS agar plates and stratified for five days. After five days, the seeds were moved to a growth room and grown under a 16-/8-h light/dark cycle at 23**°**C for 8-10 days. The seedlings were then used for TRV delivery.

A subset of transgenic T1 plants, and plants that underwent agroflood, were subjected to a heat shock treatment modified from LeBlanc et al ^31^. Seedlings that passed selection, or underwent agroflood, were then transplanted to soil and grown in a greenhouse (23°C) for one week. After one week, plants that did not receive a heat shock treatment continued to grow in the greenhouse (23°C); however, plants that underwent heat shock treatment were exposed to 8 hours (9am -5pm) of heat exposure at 37°C every day for 5 days, followed by 2 days of recovery a greenhouse (23°C). This heat shock regime lasted for two weeks.

### Protoplast isolation and transfection

Arabidopsis mesophyll protoplast isolation was performed as previously described ^20^. Plasmid transfections into *Arabidopsis* protoplasts were performed using 20µg of plasmid, similarly as ^32^. The concentrations of plasmids were determined by nanodrop. Plasmids were added to the bottom of each transfection tube, and the volume of the plasmids was supplemented with water to reach 20 µL. 200 µL of protoplasts were added followed by 220 µL of fresh and sterile polyethylene glycol (PEG)-CaCl2 solution. The samples were mixed by gently tapping the tubes, and incubated at room temperature for 10 minutes. After 10 minutes, 880 µL of W5 solution was added and mixed with the protoplasts by inverting the tube two to three times to stop the transfection. Next, protoplasts were harvested by centrifuging the tubes at 100 relative centrifugal force (RCF) for 2 min and resuspended in 1 mL of WI solution. The protoplast cells were then plated in 6-well plates pre-coated with 5% calf serum. Protoplast cells in the 6-well plates were incubated at 26°C for 48 hours. During the 48 hour incubation, the protoplast cells were subjected to a 37°C heat shock treatment for 2 hours at 16 hours post transfection. At 48 hours post transfection, protoplasts were harvested for genomic DNA extraction.

### TRV delivery to *Arabidopsis* seedlings

TRV delivery was performed as previously described ^24^. TRV1 and TRV2 vectors were first introduced into the GV3101 agrobacterium strain. The agrobacterium harboring TRV vectors were then grown in 200 ml of lysogeny broth (LB) with antibiotics for 18 hours at 28°C. Agrobacterium cultures were centrifuged for twenty minutes at 3500 RCF. The LB was discarded and the agrobacterium cells were resuspended in 200 ml of sterile water. The resuspended agrobacterium was centrifuged for 10 minutes at 3500 RPM. The supernatant was discarded and the pellet was resuspended in sterile agro-infiltration buffer containing 10 mM MgCl2, 10 mM 2-(N-morpholino) ethanesulfonic acid, and 250 µM acetosyringone to OD600 = 1.5. The agrobacterium cells were then incubated at 23°C for three hours with slow shaking. After three hours, the agrobacterium harboring TRV1 and TRV2 were mixed in a 1:1 ratio. 15 ml of the 1:1 ratio of TRV was delivered to seedlings at 8-10 days old. After four days of agroflood co-culture, seedlings were transplanted to soil.

### Screening the progeny of TRV-infected plants for edits

Seeds were harvested from the TRV-infected plants about 12 weeks after TRV delivery. The seeds were sown on ½ MS plates supplemented with 3% sucrose and stored at 4**°**C in the dark for five days to stratify. After five days, the seeds were moved to a growth room and grown under a 16-/8-h light/dark cycle at 23**°**C for 10-12 days. Amp-seq or Sanger sequencing was then performed on a subset of the progeny plants using primers listed in Supplementary Table 9.

### Next generation amplicon sequencing

DNA was extracted from protoplast samples with Qiagen DNeasy plant mini kit (Qiagen 69106). Tissue was collected from transgenic plants by sampling and pooling leaf tissue from three random leaves on a single plant three weeks after being transplanted to soil. For the plants that underwent agroflood, leaf tissue was sampled by collecting and pooling tissue from three random (however, if white sectors visible they were sampled) leaves on a single plant distal to the TRV delivery site three weeks after being transplanted to soil. Once tissue samples were collected, they were frozen at -80C overnight. The samples were then ground and DNA was extracted using the invitrogen Platinum Direct PCR Universal Master Mix (refA44647500) according to the manufacturer’s instructions. For the progenies of plants that underwent agroflood, a single leaf tissue was sampled and DNA was extracted using invitrogen Platinum Direct PCR Universal Master Mix (REF A44647500) according to the manufacturer’s instructions. The DNA was then used for Next Generation Amplicon Sequencing.

As similarly performed by Li et al ^32^, editing efficiency was characterized using single-end Next Generation Sequencing on the Illumina NovaSeqX platform. Libraries were prepared via a 2-step PCR amplification method. In the first round of amplification, each target site was amplified using primers flanking the target site (Supplementary Table 9). After 25 cycles of amplification, the reactions were cleaned using a 1.0X Ampure XP bead purification (Beckman Coulter A63881). Next, each sample went through 12 additional cycles of amplification using Illumina indexing primers. The samples were cleaned using a 0.7X Ampure XP purification. Samples were checked for purity on a 2% agarose gel, quantified using Nanodrop, normalized, and pooled.

### Next generation amplicon sequencing analysis

Amplicon sequencing analysis was performed similarly as Li et al ^32^. Only single-end reads were used for analysis. Reads were adapter trimmed using Trim Galore default settings. Remaining reads were mapped to the target genome region using the BWA aligner (v0.7.17, BWA-MEM algorithm). Sorted and indexed bam files were used as input files for further analysis by the CrispRvariants R package (v1.14.0). Each mutation pattern with corresponding read counts was exported by the CrispRvariants R package. After assessing all control samples, a criterion to classify reads as edited was established: only reads with a >= 3 bp deletion or insertion (indel) of the same pattern (indels of same size starting at the same location) with >=10 read counts from a sample were counted as edited reads. SNVs were also filtered out.

### Off-target analysis

Off-target analysis was performed as previously described in Li et al^32^. DNA from single Arabidopsis seedlings was extracted with the Qiagen DNeasy plant mini kit and sheared to 300 bp size with a Covaris. Library preparation was performed with Tecan Ovation Ultralow V2 DNA-seq kit. For variant calling, WGS reads were aligned to the TARI10 reference genome using BWA mem (v0.7.17)^33^ with default parameters. GATK (4.2.0.0)^34^ MarkDuplicatesSpark was used to remove PCR duplicate reads. Then GATK HaplotypeCaller was used to call raw variants. Raw SNPs were filtered with QD < 2.0, FS > 60.0, MQ < 40.0, and SOR > 4.0. Raw InDels were filtered with QD < 2.0, FS > 200.0, and SOR > 10.0 and used for base quality score recalibration. The recalibrated bam was further applied to GATK and Strelka (v2.9.2) SNPs/InDel calling. Only SNPs/InDels called by both GATK and Strelka were used for further filtering. The intersection of SNPs/InDel called by GATK with Strelka (v2.9.2)^35^ is obtained by BedTools (v2.26.0) ^36^. SNPs/InDel filtered with wild type background were conducted by BedTools (v2.26.0). Variants with coverage lower than 30 depth were filtered.

### RT-PCR

Total RNA from TRV-infected progeny plants was extracted using Zymo Research Direct-zol RNA MiniPrep kit (catR2052). Total RNA was converted to cDNA using the invitrogen SuperScript IV VILO Master Mix (cat11766050). The RT-PCR control was performed using primers targeting the *AtIPP2* gene (Supplementary Table 10). PCR was performed to check for the presence/absence of the TRV vector using SP9238 and SP9239 (Supplementary Table 10) ^24^. PCR was performed with New England Biolabs Q5 High-Fidelity 2X Master Mix (catM0492L) according to the manufacturer’s instructions, using 2 µl of cDNA in a 25 µl reaction. PCR conditions included a 98**°**C initial denaturation step for 30 seconds, 35x(98**°**C, 10 sec; 55**°**C, 20 sec; 72**°**C, 10 sec), 72**°**C for 2 min. 10 µl of PCR amplicons were analyzed by 2% agarose gel electrophoresis.

## Data availability

All the amp-seq data generated in this study will be accessible at NCBI Sequence Read Archive under BioProject PRJNA1124592. Whole genome sequencing data is accessible at BioProject PRJNA1146711

## Acknowledgements

This work was supported by an NSF Plant Genome Research Program grant to SEJ, JAD and JFB. SEJ and JAD are Investigators of the Howard Hughes Medical Institute. H.S. is an HHMI Fellow of The Jane Coffin Childs Fund for Medical Research. B.A.A. is supported by m-CAFEs Microbial Community Analysis & Functional Evaluation in Soils, a Science Focus Area led by Lawrence Berkeley National Laboratory based upon work supported by the US Department of Energy, Office of Science, Office of Biological & Environmental Research (DE-AC02-05CH11231). We thank Suhua Feng, Mahnaz Akhavan and the Broad Stem Cell Research Center Biosequencing Core for sequencing support, Colette Picard for assistance with analysis of amplicon sequencing data, Dinesh Kumar for supplying the TRV2 plasmid pDK3888, Feng Zhang for providing the *ku70* (SALK123114) mutant, and all members of the Jacobsen lab for their helpful insight and suggestions.

## Author Contributions

TW, MK, HS, ZL, BAA, JB, JAD and SEJ designed the research; TW, HS, ZZ, JAD, and SEJ interpreted the data; TW and SEJ wrote the manuscript; TW, MK, HS, ZL, JA, MS, KV, GW, SC, CA, NS, AS, and DS performed experiments.

---

**Extended Data Fig. 1:**
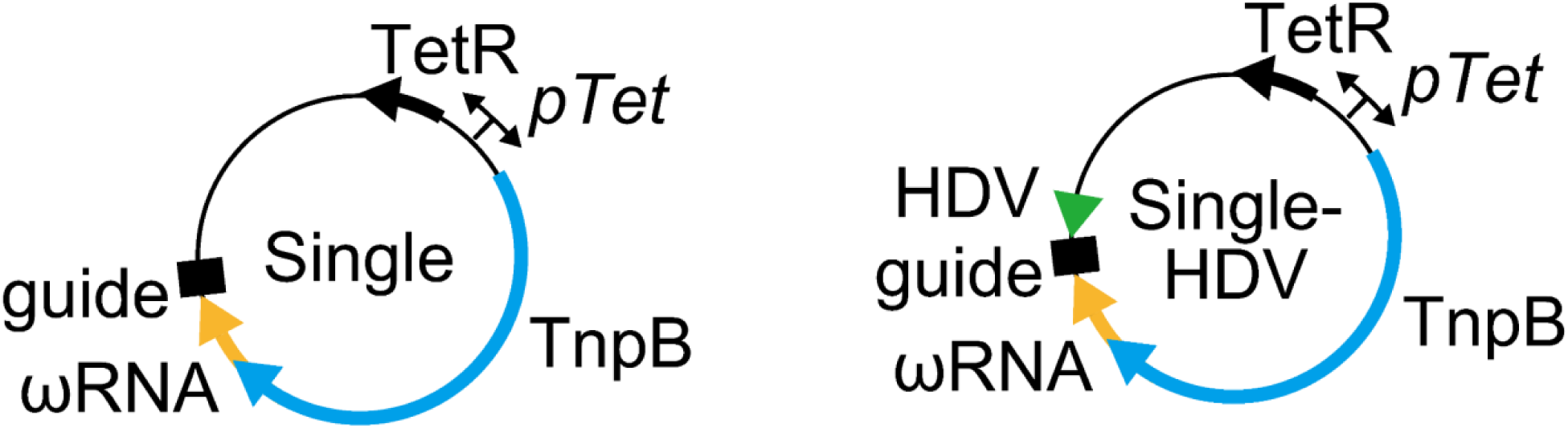
Schematic of interference assay plasmids containing either the TnpB-ωRNA Single or TnpB-ωRNA Single-HDV design. The blue arrow indicates the TnpB sequence; the yellow arrow indicates the ωRNA sequence; the black rectangle indicates the guide sequence; the green arrow indicates the HDV ribozyme sequence. The plasmids contain the tetracycline resistance gene (TetR). A tetracycline promoter (pTet) was used to drive expression of the TnpB-ωRNA single or Single-HDV sequences, and the tetracycline resistance gene.

**Extended Data Fig. 2:**
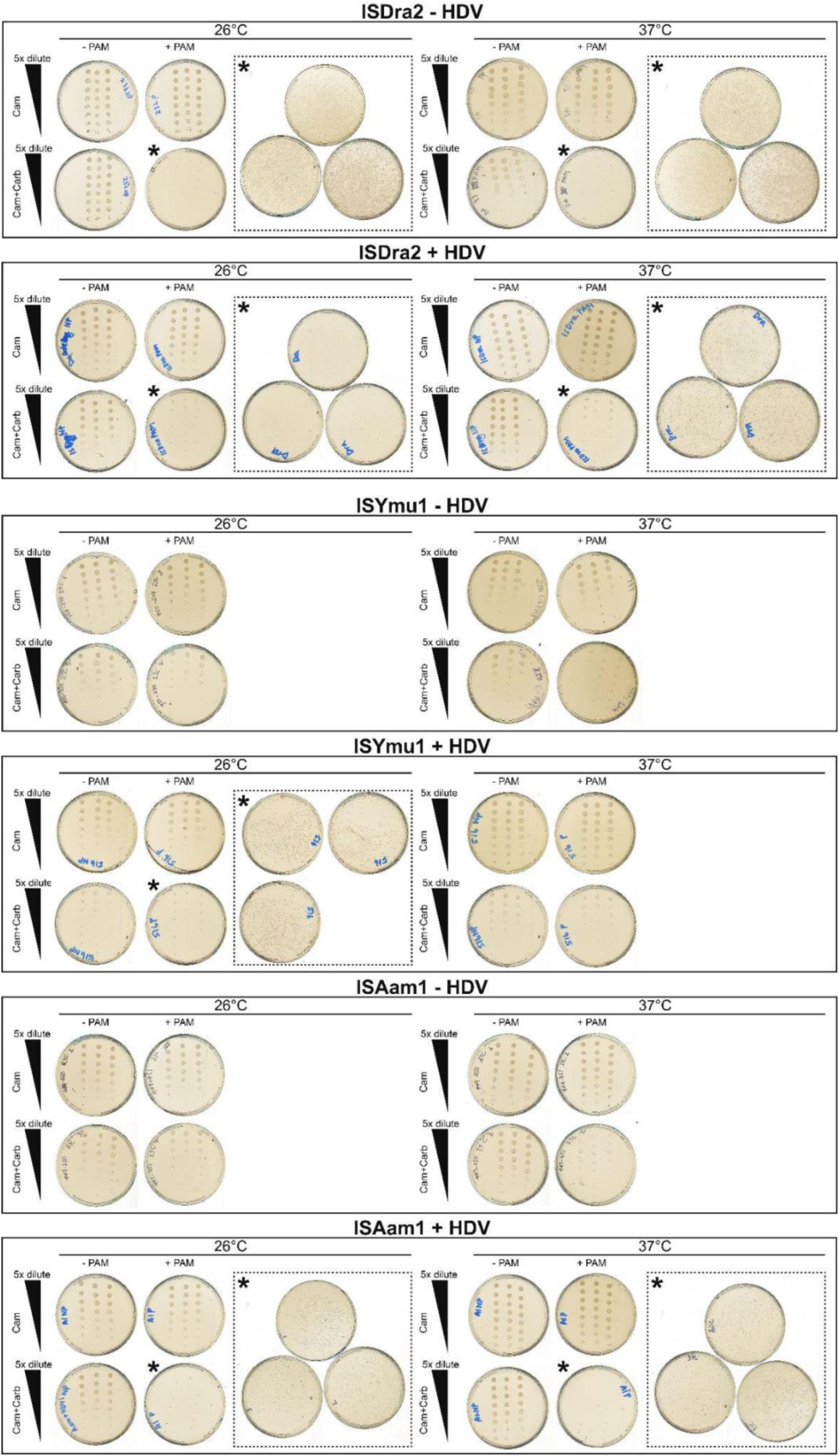
Plate images of bacterial plasmid interference assay. Five-fold serial dilutions (5 μL) from the 1 mL recovery culture post transformation were plated on both single antibiotic LB-Agar plates (Cam, upper row) and double antibiotic LB-Agar plates (Cam+Carb, lower row). Plates without visible colonies (indicated by an asterisk) had 400 μL of the original 1 mL recovery culture plated on the double antibiotic plates (dash insets). Experiments were performed in triplicates.

**Extended Data Fig. 3:**
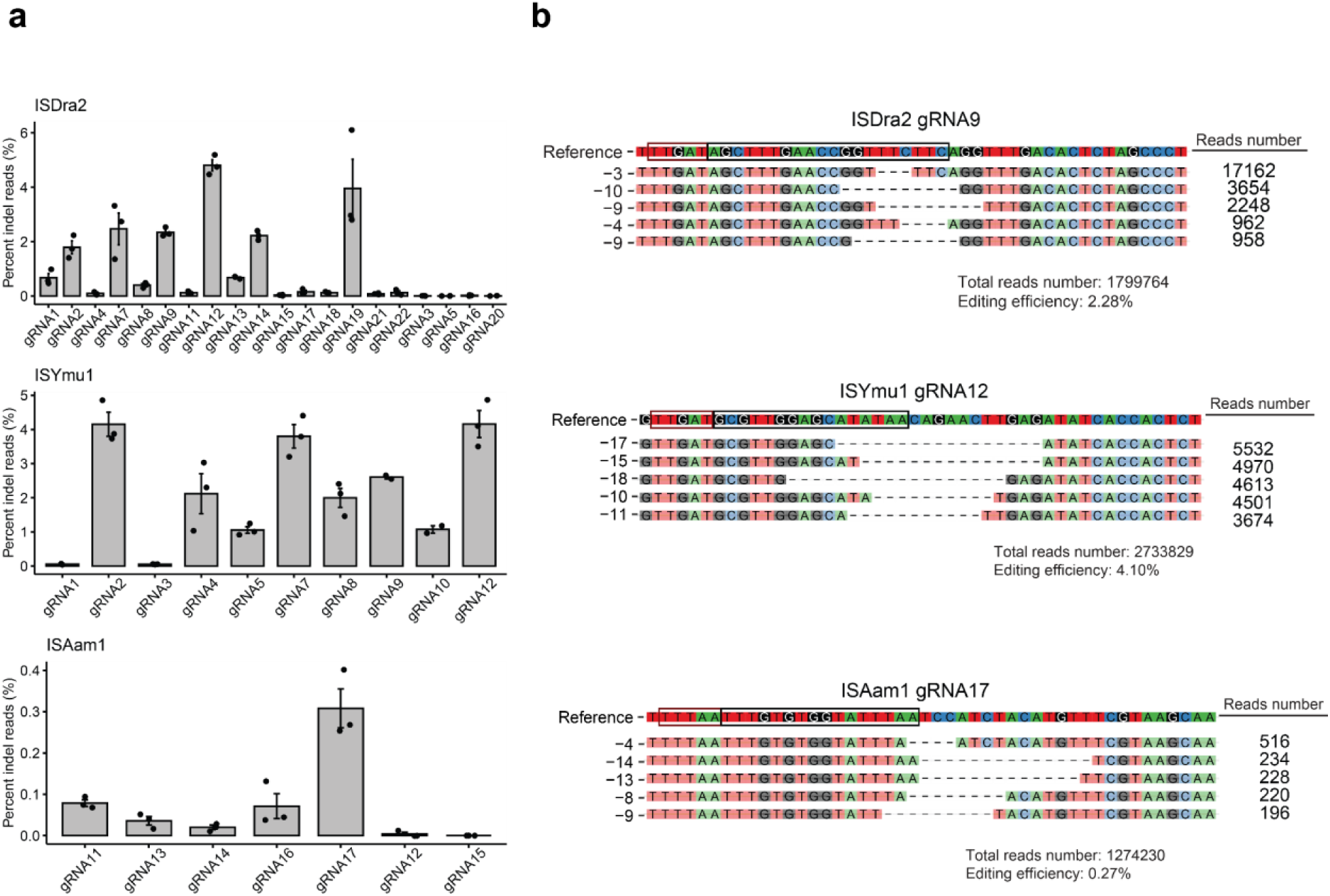
Editing efficiency and repair profiles for ISDra2, ISYmu1 and ISAam1 in protoplast experiment. **(a)** The name of each TnpB tested is at the upper left of each barplot. The gRNAs are plotted along the X-axis and the editing efficiency (percent indel reads (%)) is plotted on the Y-axis. Each dot indicates a single transfection. The standard error of the mean (SEM) was calculated for each target site. **(b)** DNA repair indel profiles for individual transfection samples. The top five most common indel types are listed on the left. The read counts for each indel are listed on the right. The PAM is identified by the red box, and the target site is outlined by the black box, in the Reference sequence. The total read number and editing efficiency are listed below each indel profile. The name of each TnpB and gRNA is displayed above the reference sequence of each indel repair profile.

**Extended Data Fig. 4:**
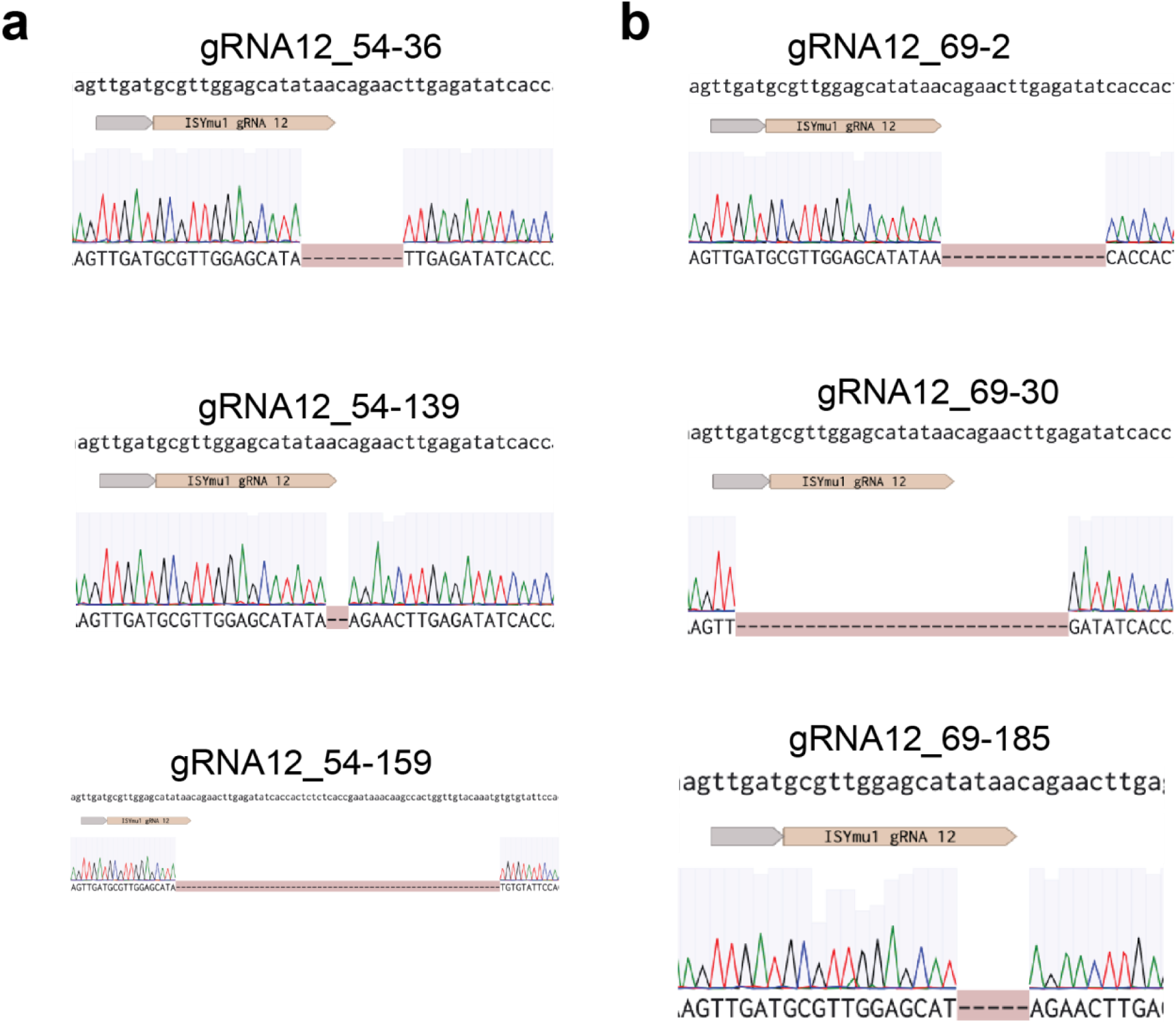
Sanger sequencing screenshots of the progeny plant genotypes for ISYmu1 gRNA12. **(a-b)** Panels a and b correspond to the progeny from plants 54 and 69, respectively. The sequence at the top is the wild type genomic sequence; below that are the ISYmu1 gRNA12 target and PAM (gray box); the ab1 trace file displays the mutation.

**Extended Data Fig. 5:**
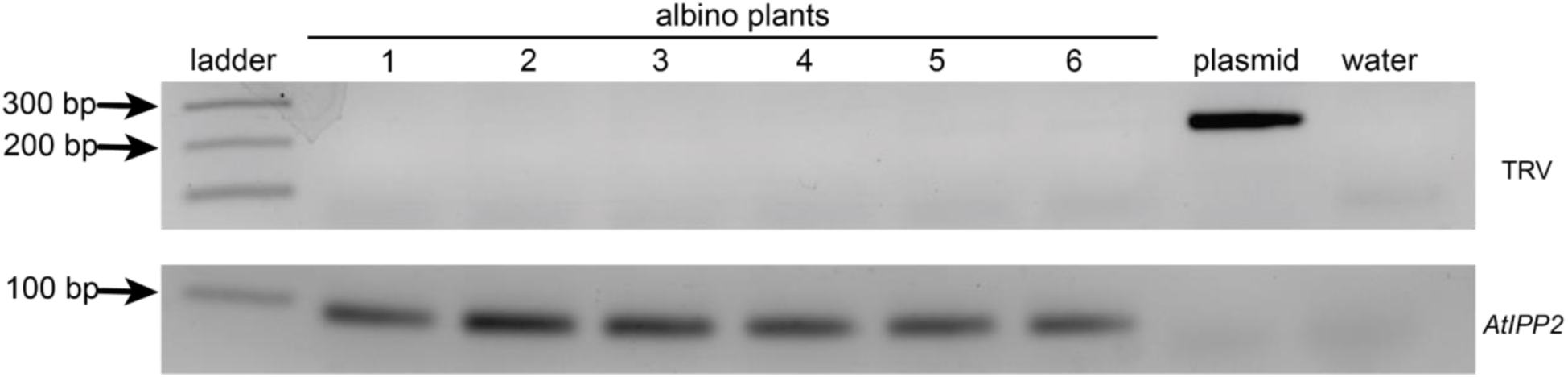
RT-PCR gel showing absence of the TRV in the progeny of a TRV-infected plant. RT-PCR was performed using total RNA extracted from albino plants homozygous for a 4bp deletion at the *AtPDS3* gene. Gel electrophoresis image for RT-PCR performed using primers targeting TRV (upper panel) and *AtIPP2* gene (bottom panel). The lanes are indicated (from left to right) as ladder, six individual albino plants, plasmid control, and a water control. Black arrows indicate the amplicon size in base pairs (bp).

**Supplementary Table 1:**
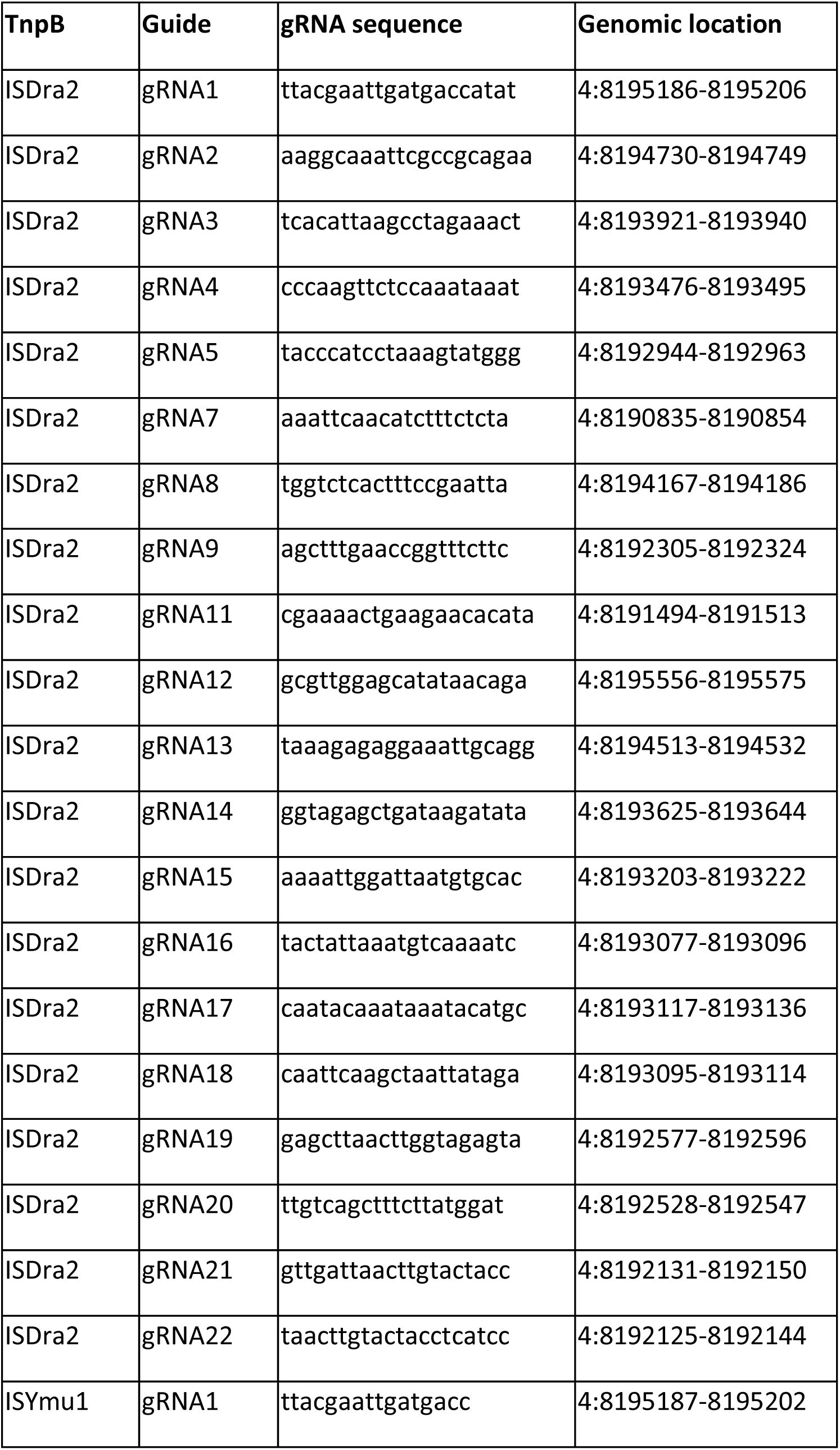
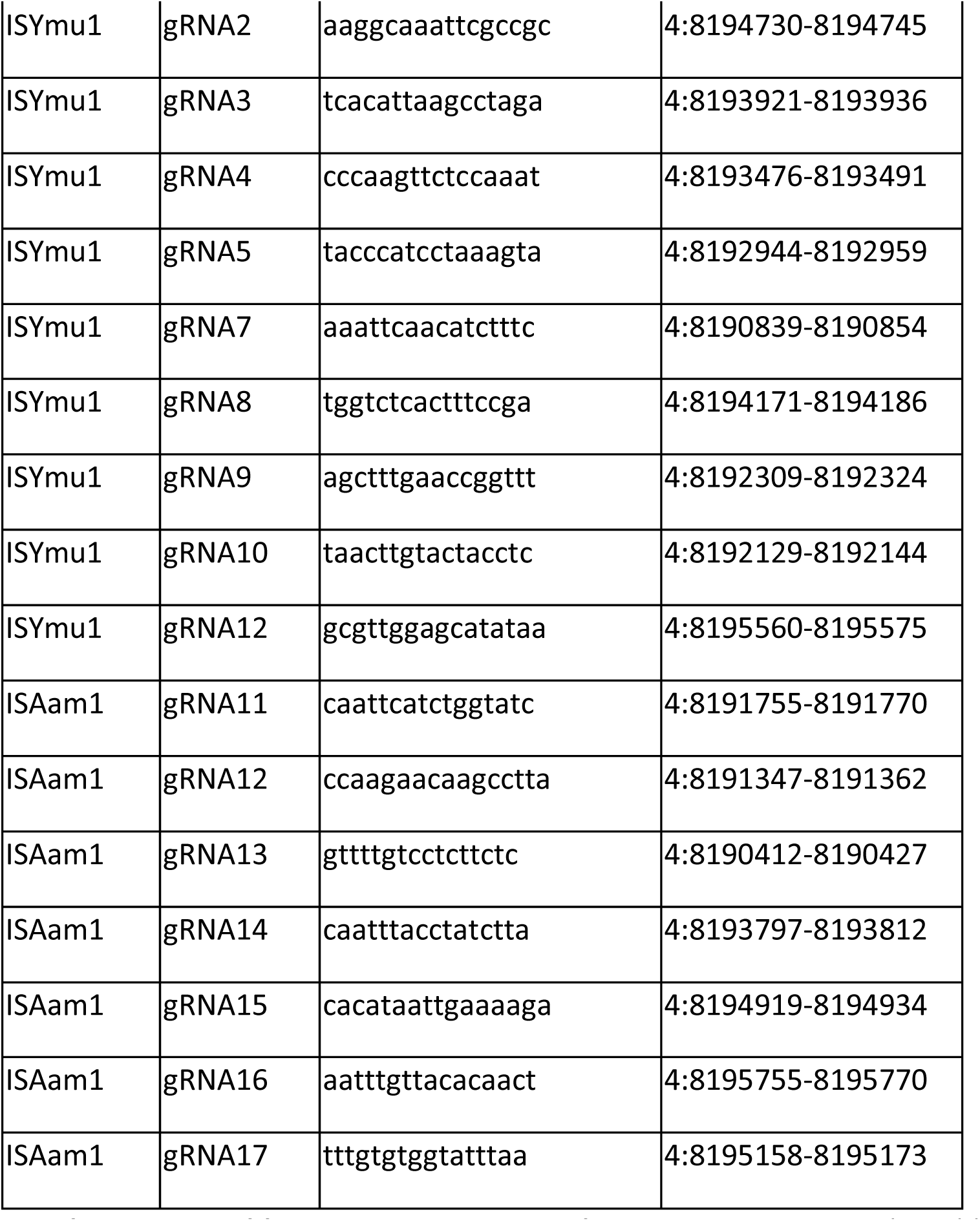
ISDra2, ISYmu1 and ISAam1 target sites. The table consists of four columns. The TnpB and Guide columns indicate the TnpB gRNA used. The gRNA sequence column lists the gRNA sequence used for targeted genome editing. The Genomic location column provides the location of each gRNA target site in the *Arabidopsis* genome.

**Supplementary Table 2:** Phenotype and genotype of the progenies from three individual TRV-infected plants. The plant ID column indicates an internal identifier for each plant sampled. The Phenotype column indicates whether the plant was green (WT) or albino. The Genotype (allele1 / allele2) column lists the alleles of the *AtPDS3* gene determined by amp-seq or Sanger Sequencing. WT is an abbreviation for wild type, bp is an abbreviation for base pair, and del is an abbreviation for deletion.

| Plant ID | Phenotype | Genotype (allele1 / allele2) | Sequencing type |
| --- | --- | --- | --- |
| gRNA2_HS_116-1 | Albino | 4bp del / 4bp del | amp-seq |
| gRNA2_HS_116-2 | Albino | 4bp del / 4bp del | amp-seq |
| gRNA2_HS_116-3 | WT | WT / WT | amp-seq |
| gRNA2_HS_116-4 | WT | WT / WT | amp-seq |
| gRNA2_HS_116-5 | WT | 4bp del / WT | amp-seq |
| gRNA2_HS_116-6 | WT | WT / WT | amp-seq |
| gRNA2_HS_116-9 | WT | WT / WT | amp-seq |
| gRNA2_HS_116-10 | WT | WT / WT | amp-seq |
| gRNA2_HS_116-12 | WT | WT / WT | amp-seq |
| gRNA2_HS_116-13 | WT | WT / WT | amp-seq |
| gRNA2_HS_116-14 | WT | WT / WT | amp-seq |
| gRNA2_HS_116-15 | WT | WT / WT | amp-seq |
| gRNA2_HS_116-16 | WT | WT / WT | amp-seq |
| gRNA2_HS_116-17 | WT | WT / WT | amp-seq |
| gRNA2_HS_116-18 | WT | WT / WT | amp-seq |
| gRNA2_HS_116-19 | WT | WT / WT | amp-seq |
| gRNA2_HS_116-20 | WT | WT / WT | amp-seq |
| gRNA2_HS_116-21 | WT | WT / WT | amp-seq |
| gRNA2_HS_116-22 | WT | WT / WT | amp-seq |
| gRNA2_HS_116-23 | WT | WT / WT | amp-seq |
| gRNA2_HS_116-24 | WT | WT / WT | amp-seq |
| gRNA2_HS_116-25 | WT | WT / WT | amp-seq |
| gRNA2_HS_116-26 | WT | WT / WT | amp-seq |
| gRNA2_HS_116-27 | WT | WT / WT | amp-seq |
| gRNA2_HS_116-28 | WT | 4bp del / WT | amp-seq |
| gRNA2_HS_116-29 | WT | WT / WT | amp-seq |
| gRNA2_HS_116-30 | Albino | 4bp del / 4bp del | amp-seq |
| gRNA2_HS_116-31 | Albino | 4bp del / 4bp del | amp-seq |
| gRNA2_HS_116-32 | Albino | 4bp del / 4bp del | amp-seq |
| gRNA2_HS_116-33 | Albino | 4bp del / 4bp del | amp-seq |
| gRNA2_HS_116-34 | Albino | 4bp del / 4bp del | amp-seq |
| gRNA2_HS_116-35 | Albino | 4bp del / 4bp del | amp-seq |
| gRNA2_HS_116-36 | Albino | 4bp del / 4bp del | amp-seq |
| gRNA2_HS_116-38 | Albino | 4bp del / 4bp del | amp-seq |
| gRNA2_HS_116-39 | Albino | 4bp del / 4bp del | amp-seq |
| gRNA2_HS_116-40 | Albino | 4bp del / 4bp del | amp-seq |
| gRNA2_HS_116-41 | Albino | 4bp del / 4bp del | amp-seq |
| gRNA2_HS_116-42 | Albino | 4bp del / 4bp del | amp-seq |
| gRNA2_HS_116-43 | Albino | 11 bp del / 17bp del | amp-seq |
| gRNA2_HS_116-45 | Albino | 4bp del / 4bp del | amp-seq |
| gRNA2_HS_116-46 | Albino | 4bp del / 4bp del | amp-seq |
| gRNA2_HS_116-47 | Albino | 4bp del / 4bp del | amp-seq |
| gRNA2_HS_116-48 | Albino | 4bp del / 4bp del | amp-seq |
| gRNA2_HS_116-49 | Albino | 10bp del / 12 bp del | amp-seq |
| gRNA2_HS_116-50 | Albino | 4bp del / 4bp del | amp-seq |
| gRNA2_HS_116-51 | Albino | 4bp del / 4bp del | amp-seq |
| gRNA2_HS_116-52 | Albino | 4bp del / 4bp del | amp-seq |
| gRNA2_HS_116-53 | Albino | 4bp del / 4bp del | amp-seq |
| gRNA2_HS_116-54 | Albino | 4bp del / 4bp del | amp-seq |
| gRNA2_HS_116-56 | Albino | 4bp del / 4bp del | amp-seq |
| gRNA2_HS_116-57 | Albino | 4bp del / 4bp del | amp-seq |
| gRNA2_HS_116-58 | Albino | 4bp del / 4bp del | amp-seq |
| gRNA2_HS_116-59 | Albino | 4bp del / 4bp del | amp-seq |
| gRNA2_HS_116-60 | Albino | 4bp del / 4bp del | amp-seq |
| gRNA2_HS_116-61 | Albino | 4bp del / 4bp del | amp-seq |
| gRNA2_HS_116-65 | Albino | 4bp del / 4bp del | amp-seq |
| gRNA2_HS_116-72 | Albino | 4bp del / 4bp del | amp-seq |
| gRNA2_HS_116-79 | Albino | 4bp del / 4bp del | amp-seq |
| gRNA2_HS_116-80 | Albino | 4bp del / 4bp del | amp-seq |
| gRNA2_HS_116-84 | Albino | 4bp del / 4bp del | amp-seq |
| gRNA2_HS_116-85 | Albino | 4bp del / 4bp del | amp-seq |
| gRNA2_HS_116-86 | Albino | 4bp del / 4bp del | amp-seq |
| gRNA2_HS_116-90 | Albino | 17bp del / 11bp del | amp-seq |
| gRNA2_HS_116-91 | Albino | 4bp del / 4bp del | amp-seq |
| gRNA2_HS_116-92 | Albino | 4bp del / 4bp del | amp-seq |
| gRNA2_HS_116-94 | WT | WT / WT | amp-seq |
| gRNA2_HS_116-95 | WT | WT / WT | amp-seq |
| gRNA2_HS_116-96 | WT | WT / WT | amp-seq |
| gRNA2_HS_116-97 | WT | WT / WT | amp-seq |
| gRNA2_HS_116-98 | WT | WT / WT | amp-seq |
| gRNA2_HS_116-99 | WT | WT / WT | amp-seq |
| gRNA2_HS_116-100 | WT | 4bp del / WT | amp-seq |
| gRNA2_HS_116-101 | WT | WT / WT | amp-seq |
| gRNA2_HS_116-102 | WT | WT / WT | amp-seq |
| gRNA2_HS_116-103 | WT | WT / WT | amp-seq |
| gRNA2_HS_116-104 | WT | WT / WT | amp-seq |
| gRNA2_HS_116-105 | WT | WT / WT | amp-seq |
| gRNA2_HS_116-106 | WT | WT / WT | amp-seq |
| gRNA2_HS_116-107 | WT | WT / WT | amp-seq |
| gRNA2_HS_116-108 | WT | 4bp del / WT | amp-seq |
| gRNA2_HS_116-109 | WT | WT / WT | amp-seq |
| gRNA2_HS_116-110 | WT | WT / WT | amp-seq |
| gRNA2_HS_116-111 | WT | WT / WT | amp-seq |
| gRNA2_HS_116-112 | WT | WT / WT | amp-seq |
| gRNA2_HS_116-113 | WT | WT / WT | amp-seq |
| gRNA2_HS_116-114 | WT | WT / WT | amp-seq |
| gRNA2_HS_116-115 | WT | WT / WT | amp-seq |
| gRNA2_HS_116-116 | WT | WT / WT | amp-seq |
| gRNA2_HS_116-117 | WT | WT / WT | amp-seq |
| gRNA2_HS_116-118 | WT | WT / WT | amp-seq |
| gRNA2_HS_116-119 | WT | WT / WT | amp-seq |
| gRNA2_HS_116-120 | WT | WT / WT | amp-seq |
| gRNA2_HS_116-122 | WT | WT / WT | amp-seq |
| gRNA2_HS_116-123 | WT | WT / WT | amp-seq |
| gRNA2_HS_116-124 | WT | WT / WT | amp-seq |
| gRNA2_HS_116-125 | WT | WT / WT | amp-seq |
| gRNA2_HS_116-126 | WT | WT / WT | amp-seq |
| gRNA2_HS_116-127 | WT | WT / WT | amp-seq |
| gRNA2_HS_116-129 | WT | WT / WT | amp-seq |
| gRNA2_HS_116-132 | WT | WT / WT | amp-seq |
| gRNA2_HS_116-133 | WT | WT / WT | amp-seq |
| gRNA2_HS_116-134 | WT | WT / WT | amp-seq |
| gRNA2_HS_116-135 | WT | WT / WT | amp-seq |
| gRNA2_HS_116-137 | WT | WT / WT | amp-seq |
| gRNA2_HS_116-138 | WT | WT / WT | amp-seq |
| gRNA2_HS_116-140 | WT | WT / WT | amp-seq |
| gRNA2_HS_116-141 | WT | WT / WT | amp-seq |
| gRNA2_HS_116-142 | WT | WT / WT | amp-seq |
| gRNA2_HS_116-143 | WT | WT / WT | amp-seq |
| gRNA2_HS_116-145 | WT | WT / WT | amp-seq |
| gRNA2_HS_116-147 | WT | WT / WT | amp-seq |
| gRNA2_HS_116-148 | WT | WT / WT | amp-seq |
| gRNA2_HS_116-149 | WT | WT / WT | amp-seq |
| gRNA2_HS_116-150 | WT | WT / WT | amp-seq |
| gRNA2_HS_116-151 | WT | WT / WT | amp-seq |
| gRNA2_HS_116-152 | WT | WT / WT | amp-seq |
| gRNA2_HS_116-153 | WT | WT / WT | amp-seq |
| gRNA2_HS_116-154 | WT | WT / WT | amp-seq |
| gRNA2_HS_116-156 | WT | WT / WT | amp-seq |
| gRNA2_HS_116-157 | WT | WT / WT | amp-seq |
| gRNA2_HS_116-158 | WT | WT / WT | amp-seq |
| gRNA2_HS_116-160 | WT | WT / WT | amp-seq |
| gRNA2_HS_116-161 | WT | WT / WT | amp-seq |
| gRNA2_HS_116-162 | WT | WT / WT | amp-seq |
| gRNA2_HS_116-163 | WT | WT / WT | amp-seq |
| gRNA2_HS_116-164 | WT | WT / WT | amp-seq |
| gRNA2_HS_116-165 | WT | WT / WT | amp-seq |
| gRNA2_HS_116-166 | WT | WT / WT | amp-seq |
| gRNA2_HS_116-167 | WT | WT / WT | amp-seq |
| gRNA2_HS_116-168 | WT | WT / WT | amp-seq |
| gRNA2_HS_116-169 | WT | WT / WT | amp-seq |
| gRNA2_HS_116-170 | WT | WT / WT | amp-seq |
| gRNA2_HS_116-171 | WT | WT / WT | amp-seq |
| gRNA2_HS_116-172 | WT | WT / WT | amp-seq |
| gRNA2_HS_116-173 | WT | WT / WT | amp-seq |
| gRNA2_HS_116-174 | WT | WT / WT | amp-seq |
| gRNA2_HS_116-175 | WT | WT / WT | amp-seq |
| gRNA2_HS_116-176 | WT | WT / WT | amp-seq |
| gRNA2_HS_116-177 | WT | WT / WT | amp-seq |
| gRNA2_HS_116-178 | WT | WT / WT | amp-seq |
| gRNA2_HS_116-179 | WT | WT / WT | amp-seq |
| gRNA2_HS_116-180 | WT | 4bp del / WT | amp-seq |
| gRNA2_HS_116-181 | WT | WT / WT | amp-seq |
| gRNA2_HS_116-182 | WT | WT / WT | amp-seq |
| gRNA2_HS_116-183 | WT | WT / WT | amp-seq |
| gRNA2_HS_116-184 | WT | WT / WT | amp-seq |
| gRNA2_HS_116-185 | WT | WT / WT | amp-seq |
| gRNA2_HS_116-186 | WT | WT / WT | amp-seq |
| gRNA2_HS_116-187 | WT | WT / WT | amp-seq |
| gRNA2_HS_116-188 | WT | WT / WT | amp-seq |
| gRNA2_HS_116-189 | WT | WT / WT | amp-seq |
| gRNA2_HS_116-190 | WT | WT / WT | amp-seq |
| gRNA2_HS_116-191 | WT | WT / WT | amp-seq |
| gRNA2_HS_116-192 | WT | WT / WT | amp-seq |
| gRNA2_HS_116-193 | WT | WT / WT | amp-seq |
| gRNA2_HS_116-194 | WT | WT / WT | amp-seq |
| gRNA2_HS_116-195 | WT | WT / WT | amp-seq |
| gRNA2_HS_116-196 | WT | WT / WT | amp-seq |
| gRNA2_HS_116-198 | WT | WT / WT | amp-seq |
| gRNA2_HS_116-199 | WT | WT / WT | amp-seq |
| gRNA2_HS_116-200 | WT | WT / WT | amp-seq |
| gRNA2_HS_116-201 | WT | WT / WT | amp-seq |
| gRNA2_HS_116-202 | WT | WT / WT | amp-seq |
| gRNA2_HS_116-203 | WT | WT / WT | amp-seq |
| gRNA2_HS_116-204 | WT | WT / WT | amp-seq |
| gRNA2_HS_116-205 | WT | WT / WT | amp-seq |
| gRNA2_HS_116-206 | WT | WT / WT | amp-seq |
| gRNA2_HS_116-207 | WT | WT / WT | amp-seq |
| gRNA2_HS_116-208 | WT | WT / WT | amp-seq |
| gRNA2_HS_116-209 | WT | WT / WT | amp-seq |
| gRNA2_HS_116-210 | WT | WT / WT | amp-seq |
| gRNA2_HS_116-211 | WT | WT / WT | amp-seq |
| gRNA2_HS_116-212 | WT | WT / WT | amp-seq |
| gRNA2_HS_116-213 | WT | WT / WT | amp-seq |
| gRNA2_HS_116-214 | WT | WT / WT | amp-seq |
| gRNA2_HS_116-215 | WT | 4bp del / WT | amp-seq |
| gRNA2_HS_116-216 | WT | WT / WT | amp-seq |
| gRNA2_HS_116-217 | WT | 4bp del / WT | amp-seq |
| gRNA2_HS_116-218 | WT | WT / WT | amp-seq |
| gRNA2_HS_116-219 | WT | WT / WT | amp-seq |
| gRNA2_HS_116-220 | WT | WT / WT | amp-seq |
| gRNA2_HS_116-221 | WT | WT / WT | amp-seq |
| gRNA2_HS_116-222 | WT | WT / WT | amp-seq |
| gRNA2_HS_116-223 | WT | WT / WT | amp-seq |
| gRNA2_HS_116-224 | WT | WT / WT | amp-seq |
| gRNA2_HS_116-225 | WT | WT / WT | amp-seq |
| gRNA2_HS_116-226 | WT | WT / WT | amp-seq |
| gRNA2_HS_116-227 | WT | WT / WT | amp-seq |
| gRNA2_HS_116-228 | WT | WT / WT | amp-seq |
| gRNA2_HS_116-229 | WT | WT / WT | amp-seq |
| gRNA2_HS_116-230 | WT | WT / WT | amp-seq |
| gRNA2_HS_116-231 | WT | WT / WT | amp-seq |
| gRNA2_HS_116-232 | WT | WT / WT | amp-seq |
| gRNA2_HS_116-233 | WT | WT / WT | amp-seq |
| gRNA2_HS_116-234 | WT | WT / WT | amp-seq |
| gRNA2_HS_116-235 | WT | WT / WT | amp-seq |
| gRNA2_HS_116-236 | WT | WT / WT | amp-seq |
| gRNA2_HS_116-237 | WT | WT / WT | amp-seq |
| gRNA2_HS_116-238 | WT | WT / WT | amp-seq |
| gRNA2_HS_116-239 | WT | WT / WT | amp-seq |
| gRNA2_HS_116-240 | WT | WT / WT | amp-seq |
| gRNA2_HS_116-241 | WT | WT / WT | amp-seq |
| gRNA2_HS_116-242 | WT | WT / WT | amp-seq |
| gRNA2_HS_116-243 | WT | WT / WT | amp-seq |
| gRNA2_HS_116-244 | WT | 4bp del / WT | amp-seq |
| gRNA2_HS_116-245 | WT | WT / WT | amp-seq |
| gRNA2_HS_116-246 | WT | WT / WT | amp-seq |
| gRNA2_HS_116-247 | WT | WT / WT | amp-seq |
| gRNA2_HS_116-248 | WT | WT / WT | amp-seq |
| gRNA12_room_54-35 | WT | 4bp del / 4bp del | Sanger |
| gRNA12_room_54-36 | WT | 9bp del / 9bp del | Sanger |
| gRNA12_room_54-37 | WT | 3bp del / 3bp del | Sanger |
| gRNA12_room_54-40 | WT | 9bp del / 9bp del | Sanger |
| gRNA12_room_54-42 | WT | 37bp del / WT | Sanger |
| gRNA12_room_54-43 | WT | WT / WT | Sanger |
| gRNA12_room_54-44 | WT | WT / WT | Sanger |
| gRNA12_room_54-45 | WT | WT / WT | Sanger |
| gRNA12_room_54-46 | WT | WT / WT | Sanger |
| gRNA12_room_54-47 | WT | WT / WT | Sanger |
| gRNA12_room_54-48 | WT | WT / WT | Sanger |
| gRNA12_room_54-49 | WT | 17bp del / 36bp del | Sanger |
| gRNA12_room_54-50 | WT | WT / WT | Sanger |
| gRNA12_room_54-52 | WT | WT / WT | Sanger |
| gRNA12_room_54-53 | WT | WT / WT | Sanger |
| gRNA12_room_54-54 | WT | 3bp del / WT | Sanger |
| gRNA12_room_54-55 | WT | WT / WT | Sanger |
| gRNA12_room_54-58 | WT | 17bp del / 36bp del | Sanger |
| gRNA12_room_54-59 | WT | 9bp del / 9bp del | Sanger |
| gRNA12_room_54-60 | WT | WT / WT | Sanger |
| gRNA12_room_54-61 | WT | 17bp del / 36bp del | Sanger |
| gRNA12_room_54-62 | WT | WT / WT | Sanger |
| gRNA12_room_54-63 | WT | WT / WT | Sanger |
| gRNA12_room_54-64 | WT | WT / WT | Sanger |
| gRNA12_room_54-65 | WT | WT / WT | Sanger |
| gRNA12_room_54-66 | WT | WT / WT | Sanger |
| gRNA12_room_54-68 | WT | 12bp del / 12bp del | Sanger |
| gRNA12_room_54-69 | WT | 3bp del / 3bp del | Sanger |
| gRNA12_room_54-70 | WT | WT / WT | Sanger |
| gRNA12_room_54-71 | WT | WT / WT | Sanger |
| gRNA12_room_54-72 | WT | 10bp del / WT | Sanger |
| gRNA12_room_54-73 | WT | WT / WT | Sanger |
| gRNA12_room_54-74 | WT | WT / WT | Sanger |
| gRNA12_room_54-75 | WT | WT / WT | Sanger |
| gRNA12_room_54-76 | WT | WT / WT | Sanger |
| gRNA12_room_54-77 | WT | WT / WT | Sanger |
| gRNA12_room_54-79 | WT | 4bp del / WT | Sanger |
| gRNA12_room_54-80 | WT | WT / WT | Sanger |
| gRNA12_room_54-84 | WT | 9bp del / WT | Sanger |
| gRNA12_room_54-86 | WT | WT / WT | Sanger |
| gRNA12_room_54-90 | WT | WT / WT | Sanger |
| gRNA12_room_54-91 | WT | 9bp del / 9bp del | Sanger |
| gRNA12_room_54-94 | WT | WT / WT | Sanger |
| gRNA12_room_54-95 | WT | WT / WT | Sanger |
| gRNA12_room_54-97 | WT | 3bp del / 3bp del | Sanger |
| gRNA12_room_54-98 | WT | WT / WT | Sanger |
| gRNA12_room_54-104 | WT | WT / WT | Sanger |
| gRNA12_room_54-105 | WT | 7bp del / 7bp del | Sanger |
| gRNA12_room_54-106 | WT | WT / WT | Sanger |
| gRNA12_room_54-108 | WT | WT / WT | Sanger |
| gRNA12_room_54-109 | WT | 17bp del / 21bp del | Sanger |
| gRNA12_room_54-110 | WT | 3bp del / WT | Sanger |
| gRNA12_room_54-111 | WT | WT / WT | Sanger |
| gRNA12_room_54-112 | WT | 7bp del / 7bp del | Sanger |
| gRNA12_room_54-113 | WT | 26bp del / WT | Sanger |
| gRNA12_room_54-114 | WT | WT / WT | Sanger |
| gRNA12_room_54-115 | WT | WT / WT | Sanger |
| gRNA12_room_54-116 | WT | WT / WT | Sanger |
| gRNA12_room_54-117 | WT | WT / WT | Sanger |
| gRNA12_room_54-119 | WT | WT / WT | Sanger |
| gRNA12_room_54-120 | WT | 11bp del / 11bp del | Sanger |
| gRNA12_room_54-121 | WT | 10bp del / WT | Sanger |
| gRNA12_room_54-122 | WT | WT / WT | Sanger |
| gRNA12_room_54-123 | WT | WT / WT | Sanger |
| gRNA12_room_54-124 | WT | 10bp del / WT | Sanger |
| gRNA12_room_54-125 | WT | WT / WT | Sanger |
| gRNA12_room_54-126 | WT | WT / WT | Sanger |
| gRNA12_room_54-127 | WT | WT / WT | Sanger |
| gRNA12_room_54-128 | WT | 4bp del / WT | Sanger |
| gRNA12_room_54-130 | WT | 38bp del / WT | Sanger |
| gRNA12_room_54-132 | WT | WT / WT | Sanger |
| gRNA12_room_54-133 | WT | WT / WT | Sanger |
| gRNA12_room_54-134 | WT | WT / WT | Sanger |
| gRNA12_room_54-135 | WT | WT / WT | Sanger |
| gRNA12_room_54-136 | WT | WT / WT | Sanger |
| gRNA12_room_54-138 | WT | 9bp del / 9bp del | Sanger |
| gRNA12_room_54-139 | WT | 2bp del / 2bp del | Sanger |
| gRNA12_room_54-140 | WT | WT / WT | Sanger |
| gRNA12_room_54-141 | WT | WT / WT | Sanger |
| gRNA12_room_54-142 | WT | 37bp del / WT | Sanger |
| gRNA12_room_54-143 | WT | WT / WT | Sanger |
| gRNA12_room_54-144 | WT | WT / WT | Sanger |
| gRNA12_room_54-145 | WT | WT / WT | Sanger |
| gRNA12_room_54-146 | WT | WT / WT | Sanger |
| gRNA12_room_54-148 | WT | 17bp del / 36bp del | Sanger |
| gRNA12_room_54-149 | WT | WT / WT | Sanger |
| gRNA12_room_54-152 | WT | WT / WT | Sanger |
| gRNA12_room_54-153 | WT | WT / WT | Sanger |
| gRNA12_room_54-154 | WT | WT / WT | Sanger |
| gRNA12_room_54-155 | WT | WT / WT | Sanger |
| gRNA12_room_54-156 | WT | 11bp del / 11bp del | Sanger |
| gRNA12_room_54-157 | WT | 5bp del / WT | Sanger |
| gRNA12_room_54-158 | WT | 37bp del / WT | Sanger |
| gRNA12_room_54-159 | WT | 62bp del / 62bp del | Sanger |
| gRNA12_room_54-160 | WT | WT / WT | Sanger |
| gRNA12_room_54-161 | WT | 8bp del / WT | Sanger |
| gRNA12_room_54-162 | WT | 8bp del / WT | Sanger |
| gRNA12_room_54-163 | WT | WT / WT | Sanger |
| gRNA12_room_54-164 | WT | 4bp del / WT | Sanger |
| gRNA12_room_54-165 | WT | WT / WT | Sanger |
| gRNA12_room_54-166 | WT | 2bp del / WT | Sanger |
| gRNA12_room_54-167 | WT | WT / WT | Sanger |
| gRNA12_room_54-168 | WT | WT / WT | Sanger |
| gRNA12_room_54-169 | WT | WT / WT | Sanger |
| gRNA12_room_54-170 | WT | WT / WT | Sanger |
| gRNA12_room_54-171 | WT | WT / WT | Sanger |
| gRNA12_room_54-172 | WT | WT / WT | Sanger |
| gRNA12_room_54-173 | WT | WT / WT | Sanger |
| gRNA12_room_54-175 | WT | WT / WT | Sanger |
| gRNA12_room_54-176 | WT | WT / WT | Sanger |
| gRNA12_room_54-178 | WT | WT / WT | Sanger |
| gRNA12_room_54-179 | WT | WT / WT | Sanger |
| gRNA12_room_54-180 | WT | WT / WT | Sanger |
| gRNA12_room_54-181 | WT | WT / WT | Sanger |
| gRNA12_room_54-182 | WT | WT / WT | Sanger |
| gRNA12_room_54-183 | WT | WT / WT | Sanger |
| gRNA12_room_54-184 | WT | 5bp del / WT | Sanger |
| gRNA12_room_54-185 | WT | WT / WT | Sanger |
| gRNA12_room_54-186 | WT | WT / WT | Sanger |
| gRNA12_room_54-190 | WT | 9bp del / WT | Sanger |
| gRNA12_room_54-191 | WT | 15bp del / WT | Sanger |
| gRNA12_room_54-192 | WT | WT / WT | Sanger |
| gRNA12_room_54-193 | WT | WT / WT | Sanger |
| gRNA12_room_54-194 | WT | 7bp del / 7bp del | Sanger |
| gRNA12_room_54-195 | WT | WT / WT | Sanger |
| gRNA12_room_54-196 | WT | WT / WT | Sanger |
| gRNA12_room_54-197 | WT | 10bp del / 10bp del | Sanger |
| gRNA12_room_54-198 | WT | 10bp del / WT | Sanger |
| gRNA12_room_54-199 | WT | WT / WT | Sanger |
| gRNA12_room_54-200 | WT | 2bp del / 2bp del | Sanger |
| gRNA12_room_54-202 | WT | WT / WT | Sanger |
| gRNA12_room_54-205 | WT | WT / WT | Sanger |
| gRNA12_room_54-208 | WT | WT / WT | Sanger |
| gRNA12_room_54-209 | WT | WT / WT | Sanger |
| gRNA12_room_54-211 | WT | WT / WT | Sanger |
| gRNA12_room_54-212 | WT | WT / WT | Sanger |
| gRNA12_room_54-213 | WT | 3bp del / WT | Sanger |
| gRNA12_room_54-214 | WT | WT / WT | Sanger |
| gRNA12_room_54-215 | WT | 4bp del / 4bp del | Sanger |
| gRNA12_room_54-216 | WT | WT / WT | Sanger |
| gRNA12_room_54-217 | WT | 3bp del / WT | Sanger |
| gRNA12_room_54-218 | WT | WT / WT | Sanger |
| gRNA12_room_54-219 | WT | 17bp del / 17bp del | Sanger |
| gRNA12_room_54-220 | WT | 4bp del / WT | Sanger |
| gRNA12_room_54-221 | WT | 7bp del / 10bp del | Sanger |
| gRNA12_room_54-222 | WT | WT / WT | Sanger |
| gRNA12_room_54-223 | WT | WT / WT | Sanger |
| gRNA12_room_54-224 | WT | WT / WT | Sanger |
| gRNA12_room_69-2 | WT | 15bp del / 15bp del | Sanger |
| gRNA12_room_69-4 | WT | 27bp del / WT | Sanger |
| gRNA12_room_69-5 | WT | 5bp del / 24bp del | Sanger |
| gRNA12_room_69-7 | WT | WT / WT | Sanger |
| gRNA12_room_69-8 | WT | 7bp del / 15bp del | Sanger |
| gRNA12_room_69-9 | WT | 1bp del / 1bp del | Sanger |
| gRNA12_room_69-10 | WT | WT / WT | Sanger |
| gRNA12_room_69-12 | WT | 5bp del / 15bp del | Sanger |
| gRNA12_room_69-14 | WT | 22bp del / 22bp del | Sanger |
| gRNA12_room_69-16 | WT | 10bp del / 10bp del | Sanger |
| gRNA12_room_69-17 | WT | 1bp del / 17bp del | Sanger |
| gRNA12_room_69-18 | WT | 11bp del / WT | Sanger |
| gRNA12_room_69-21 | WT | 5bp del / 24bp del | Sanger |
| gRNA12_room_69-22 | WT | 17bp del / WT | Sanger |
| gRNA12_room_69-23 | WT | WT / WT | Sanger |
| gRNA12_room_69-28 | WT | 5bp del / 4bp del | Sanger |
| gRNA12_room_69-30 | WT | 29bp del / 29bp del | Sanger |
| gRNA12_room_69-33 | WT | WT / WT | Sanger |
| gRNA12_room_69-34 | WT | 5bp del / 5bp del | Sanger |
| gRNA12_room_69-37 | WT | WT / WT | Sanger |
| gRNA12_room_69-41 | WT | WT / WT | Sanger |
| gRNA12_room_69-44 | WT | 5bp del / 15bp del | Sanger |
| gRNA12_room_69-51 | WT | 5bp del / WT | Sanger |
| gRNA12_room_69-53 | WT | 15bp del / 15bp del | Sanger |
| gRNA12_room_69-55 | WT | 5bp del / 5bp del | Sanger |
| gRNA12_room_69-59 | WT | 24bp del / 24bp del | Sanger |
| gRNA12_room_69-63 | WT | 5bp del / 24bp del | Sanger |
| gRNA12_room_69-66 | WT | WT / WT | Sanger |
| gRNA12_room_69-67 | WT | WT / WT | Sanger |
| gRNA12_room_69-70 | WT | 7bp del / WT | Sanger |
| gRNA12_room_69-73 | WT | 17bp del / WT | Sanger |
| gRNA12_room_69-74 | WT | 5bp del / 10bp del | Sanger |
| gRNA12_room_69-78 | WT | WT / WT | Sanger |
| gRNA12_room_69-80 | WT | WT / WT | Sanger |
| gRNA12_room_69-87 | WT | 13bp del / 14bp del | Sanger |
| gRNA12_room_69-88 | WT | 11bp del / WT | Sanger |
| gRNA12_room_69-91 | WT | WT / WT | Sanger |
| gRNA12_room_69-92 | WT | 5bp del / 10bp del | Sanger |
| gRNA12_room_69-93 | WT | WT / WT | Sanger |
| gRNA12_room_69-94 | WT | WT / WT | Sanger |
| gRNA12_room_69-95 | WT | 3bp del / 23bp del | Sanger |
| gRNA12_room_69-101 | WT | 7bp del / WT | Sanger |
| gRNA12_room_69-108 | WT | WT / WT | Sanger |
| gRNA12_room_69-111 | WT | WT / WT | Sanger |
| gRNA12_room_69-115 | WT | WT / WT | Sanger |
| gRNA12_room_69-117 | WT | 10bp del / WT | Sanger |
| gRNA12_room_69-120 | WT | 17bp del / WT | Sanger |
| gRNA12_room_69-123 | WT | WT / WT | Sanger |
| gRNA12_room_69-124 | WT | 15bp del / 15bp del | Sanger |
| gRNA12_room_69-125 | WT | 9bp del / 22bp del | Sanger |
| gRNA12_room_69-126 | WT | 17bp del / 17bp del | Sanger |
| gRNA12_room_69-127 | WT | WT / WT | Sanger |
| gRNA12_room_69-129 | WT | WT / WT | Sanger |
| gRNA12_room_69-130 | WT | WT / WT | Sanger |
| gRNA12_room_69-131 | WT | 4bp del / 5bp del | Sanger |
| gRNA12_room_69-132 | WT | WT / WT | Sanger |
| gRNA12_room_69-134 | WT | WT / WT | Sanger |
| gRNA12_room_69-137 | WT | 4bp del / WT | Sanger |
| gRNA12_room_69-142 | WT | 15bp del / 5bp del | Sanger |
| gRNA12_room_69-148 | WT | WT / WT | Sanger |
| gRNA12_room_69-149 | WT | 22bp del / 22bp del | Sanger |
| gRNA12_room_69-150 | WT | WT / WT | Sanger |
| gRNA12_room_69-151 | WT | WT / WT | Sanger |
| gRNA12_room_69-154 | WT | 11bp del / 11bp del | Sanger |
| gRNA12_room_69-157 | WT | WT / WT | Sanger |
| gRNA12_room_69-158 | WT | 11bp del / WT | Sanger |
| gRNA12_room_69-159 | WT | 18bp del / WT | Sanger |
| gRNA12_room_69-163 | WT | WT / WT | Sanger |
| gRNA12_room_69-170 | WT | 7bp del / 26bp del | Sanger |
| gRNA12_room_69-171 | WT | 3bp del / 5bp del | Sanger |
| gRNA12_room_69-172 | WT | 10bp del / 10bp del | Sanger |
| gRNA12_room_69-175 | WT | 5bp del / WT | Sanger |
| gRNA12_room_69-183 | WT | WT / WT | Sanger |
| gRNA12_room_69-185 | WT | 5bp del / 5bp del | Sanger |
| gRNA12_room_69-188 | WT | 1bp del / WT | Sanger |

**Supplementary Table 3:** Whole genome sequencing read count and mapping rate of WT control and albino mutant plants.

| Sample | Reads count | Unmapped reads count | Mapping rate (%) | Coverage |
| --- | --- | --- | --- | --- |
| WT-rep1 | 331996121 | 2122433 | 99.36 | 817.69 |
| WT-rep2 | 314298698 | 1969352 | 99.37 | 774.20 |
| gRNA2_HS_116-2 | 315840117 | 1628611 | 99.48 | 778.87 |
| gRNA2_HS_116-32 | 357227432 | 2397598 | 99.33 | 879.55 |
| gRNA2_HS_116-43 | 265745567 | 1688428 | 99.36 | 654.55 |

**Supplementary Table 4:** WGS variant detection of WT control and albino mutant plants using GATK and Strelka2.

| Sample | GATK | Streak2 | GATK+Streak2 | Filter with WT | Filter by depth > 30 | Manually check |
| --- | --- | --- | --- | --- | --- | --- |
| WT | 40852 | 24406 | NA | NA | NA | NA |
| albino-plant2 | 36346 | 20659 | 16296 | 28 | 25 | 5 |
| albino-plant32 | 36342 | 20991 | 16629 | 35 | 30 | 5 |
| albino-plant43 | 36989 | 21928 | 17111 | 80 | 71 | 4 |

**Supplementary Table 5:**
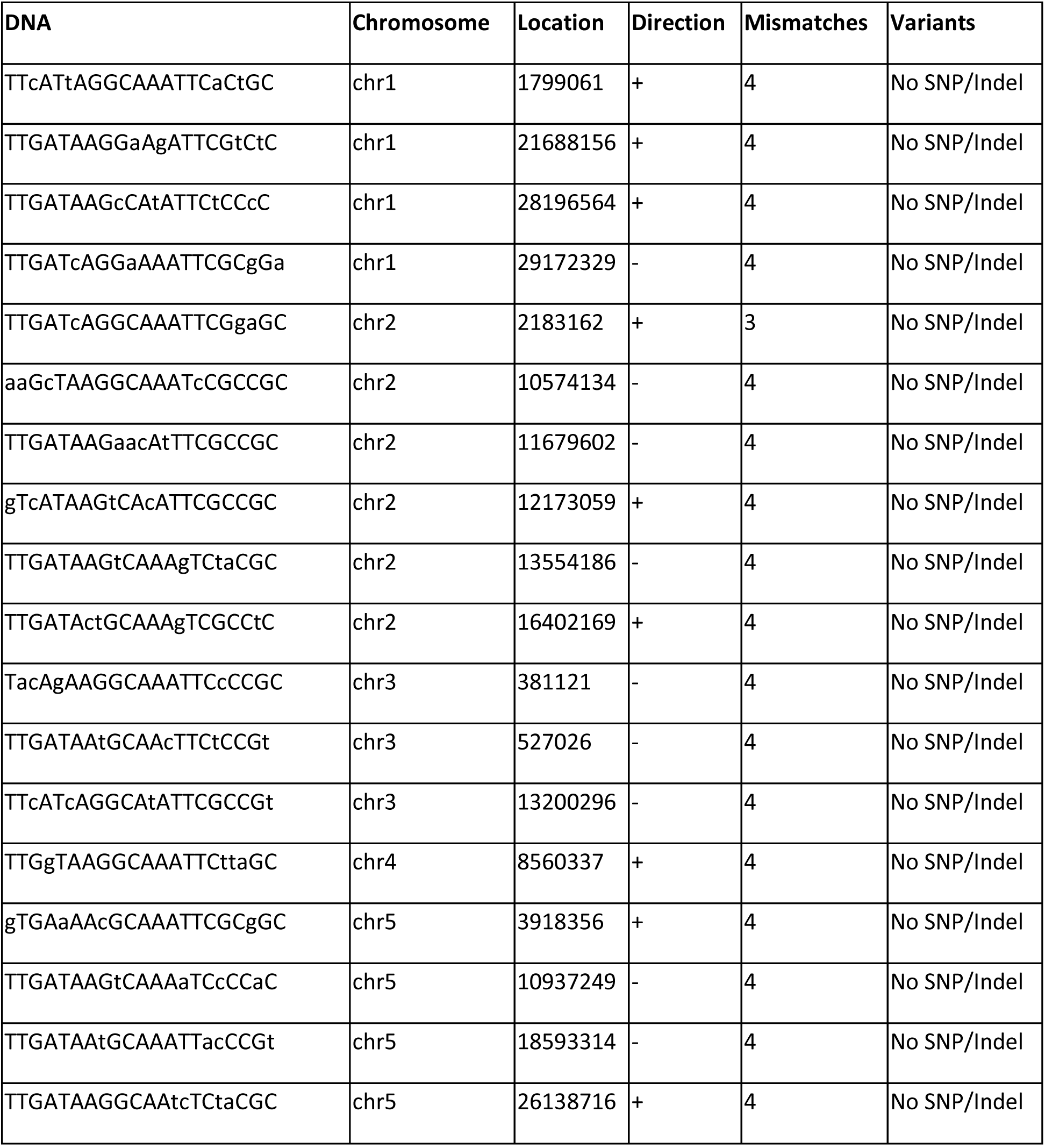
Off-target editing analysis using Cas-OFFinder. The DNA column contains the sequence of the potential off-target site, with lower case letters indicating a mismatch compared to the actual target site. The Chromosome, Location, and Direction columns indicate where in the *Arabidopsis* genome that potential off-target site is located. The mismatch column indicates the number of mismatches in the potential off-target site relative to the actual target site sequence. The Variants column lists the off-target editing result for that site, with every off-target site analyzed as wild type (No SNP/Indel).

**Supplementary Table 6:**
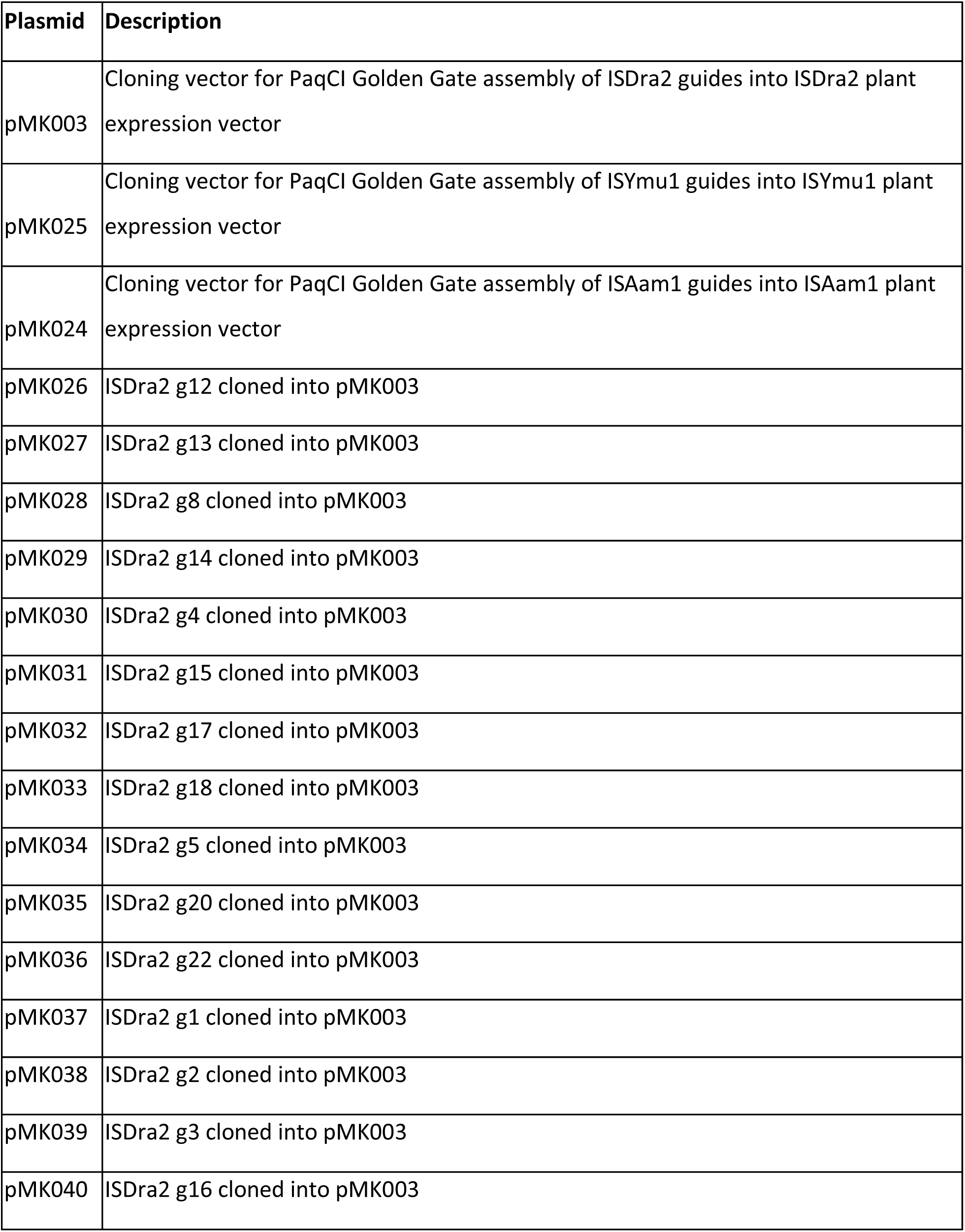
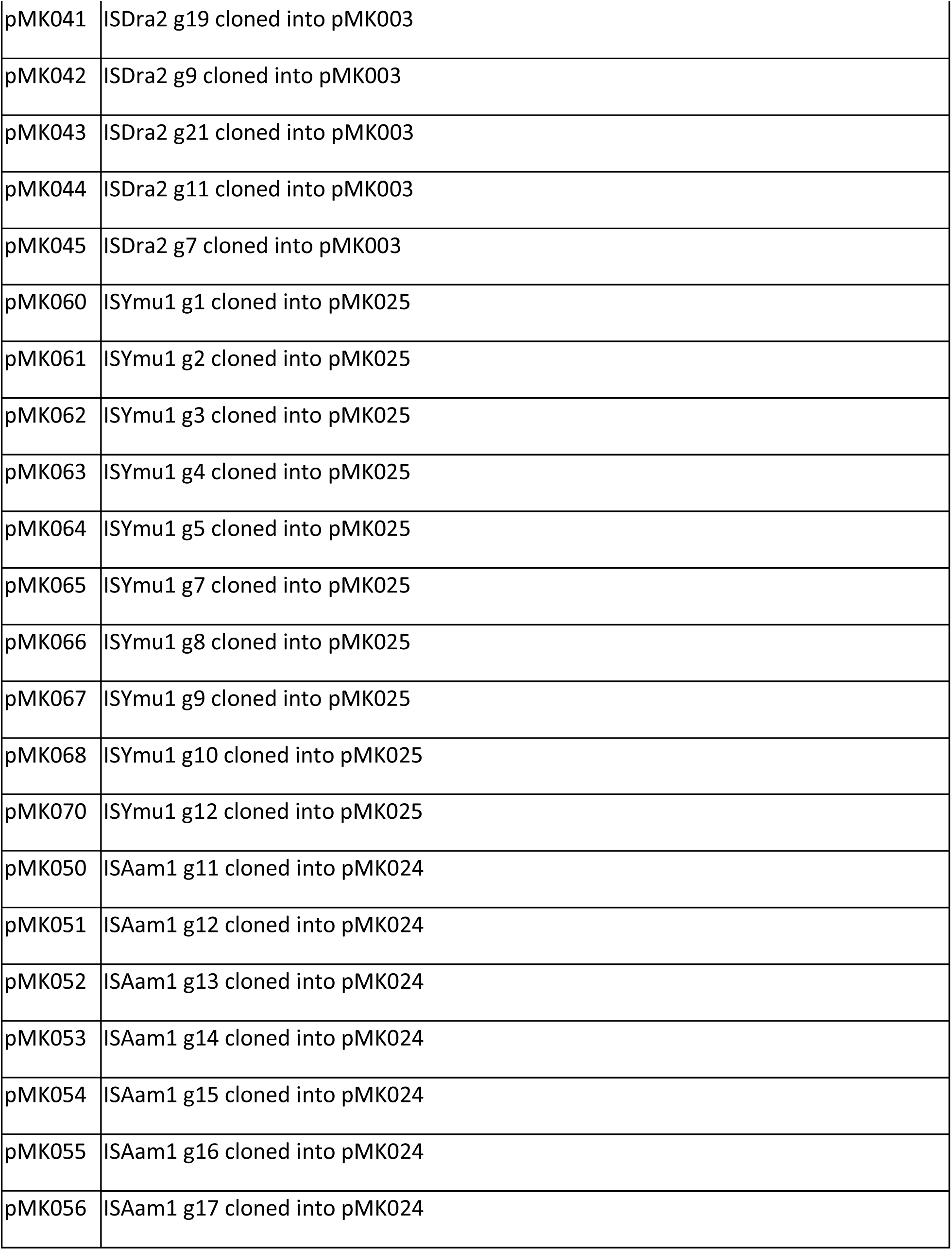
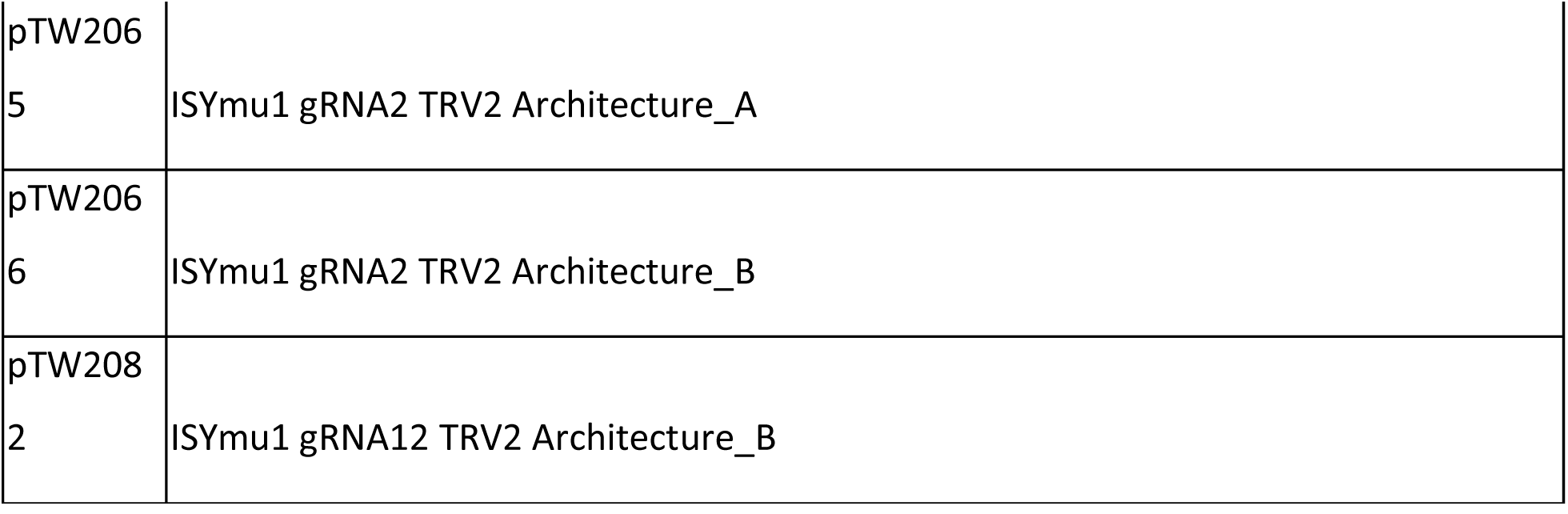
Plasmids used in this study. The Plasmid column lists the name of each plasmid used. The Description column provides a TnpB and gRNA description of each plasmid used, along with the intermediate plant expression vector that was used to create it.

**Supplementary Table 7:**
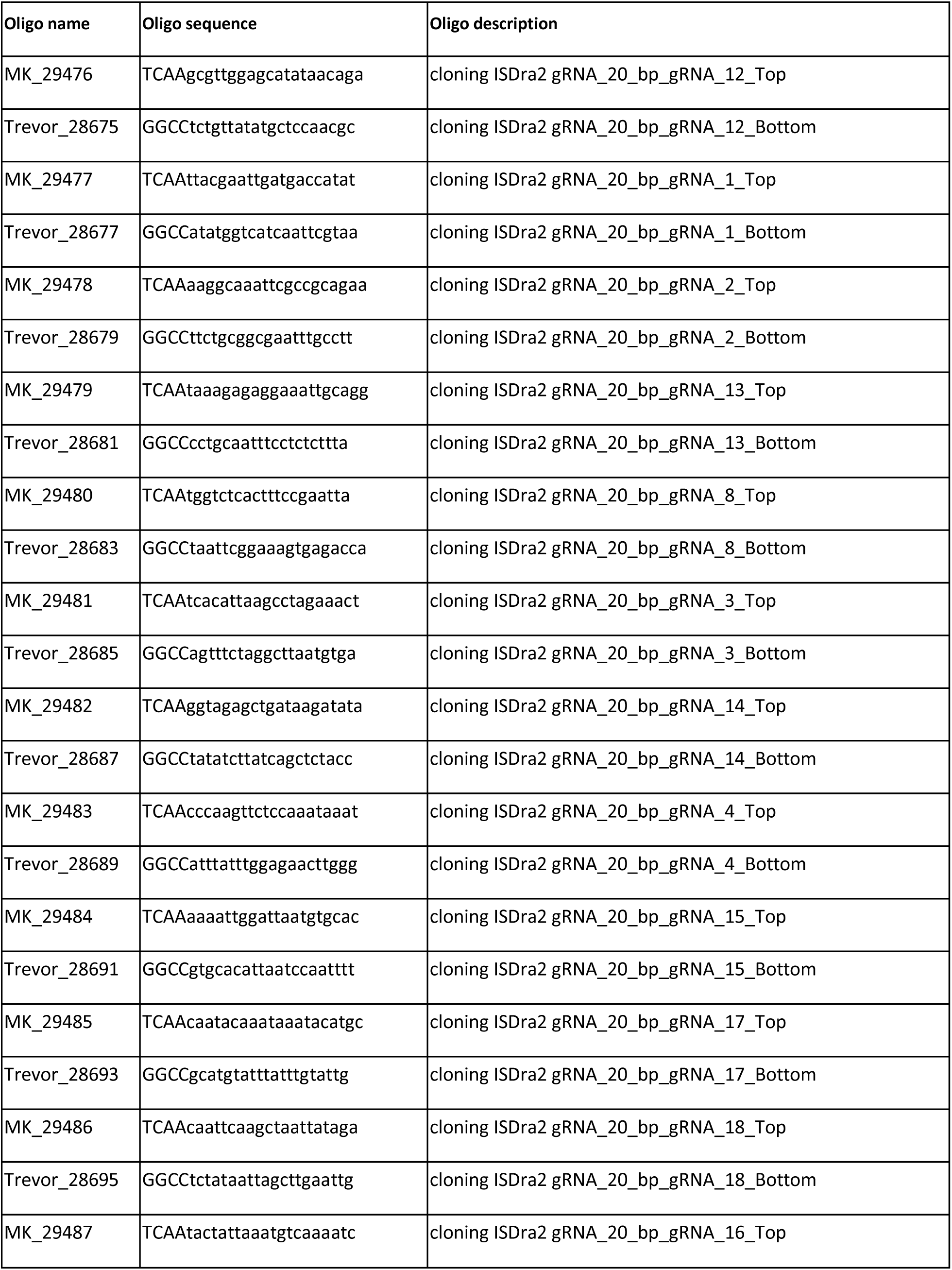
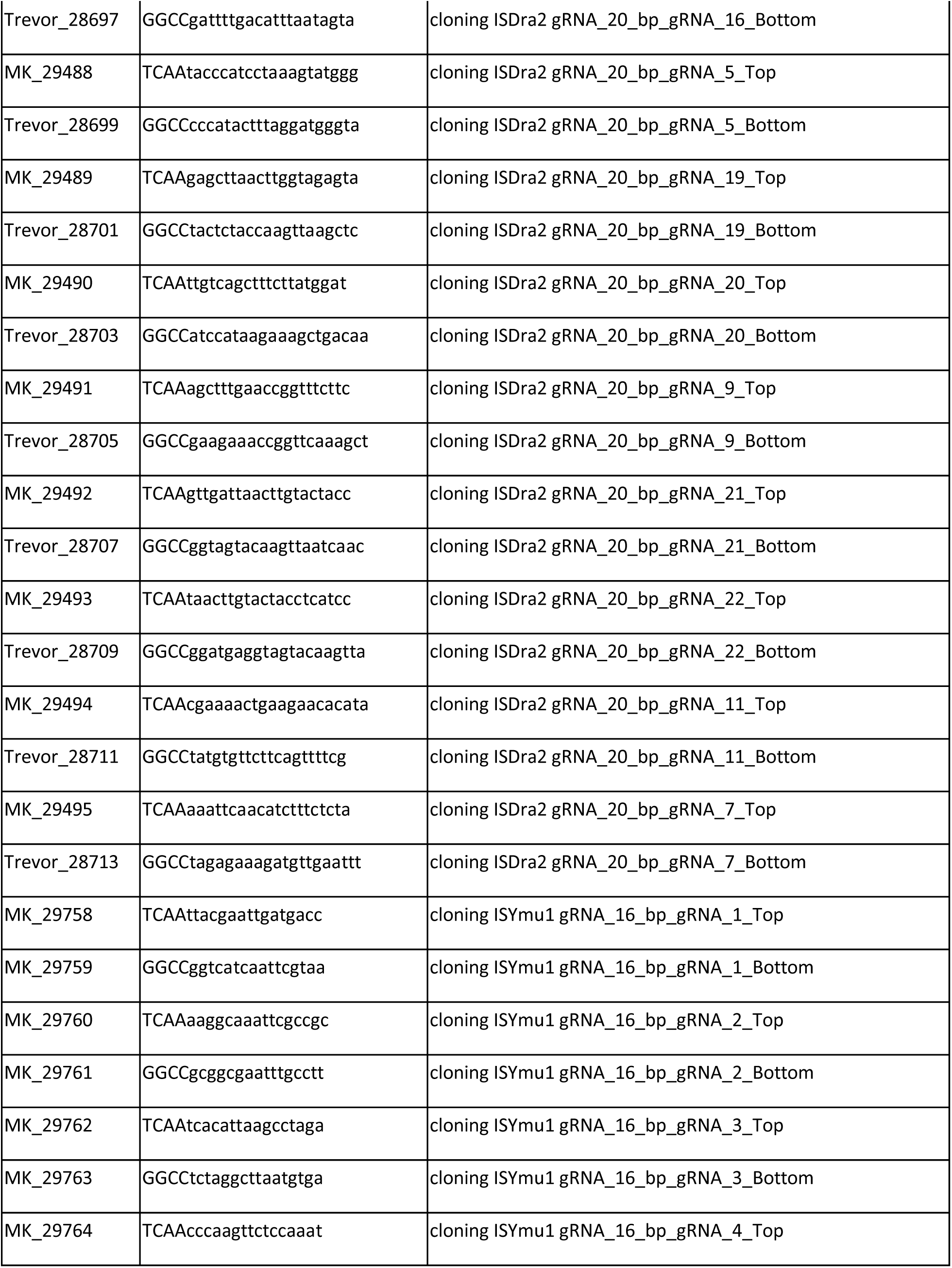
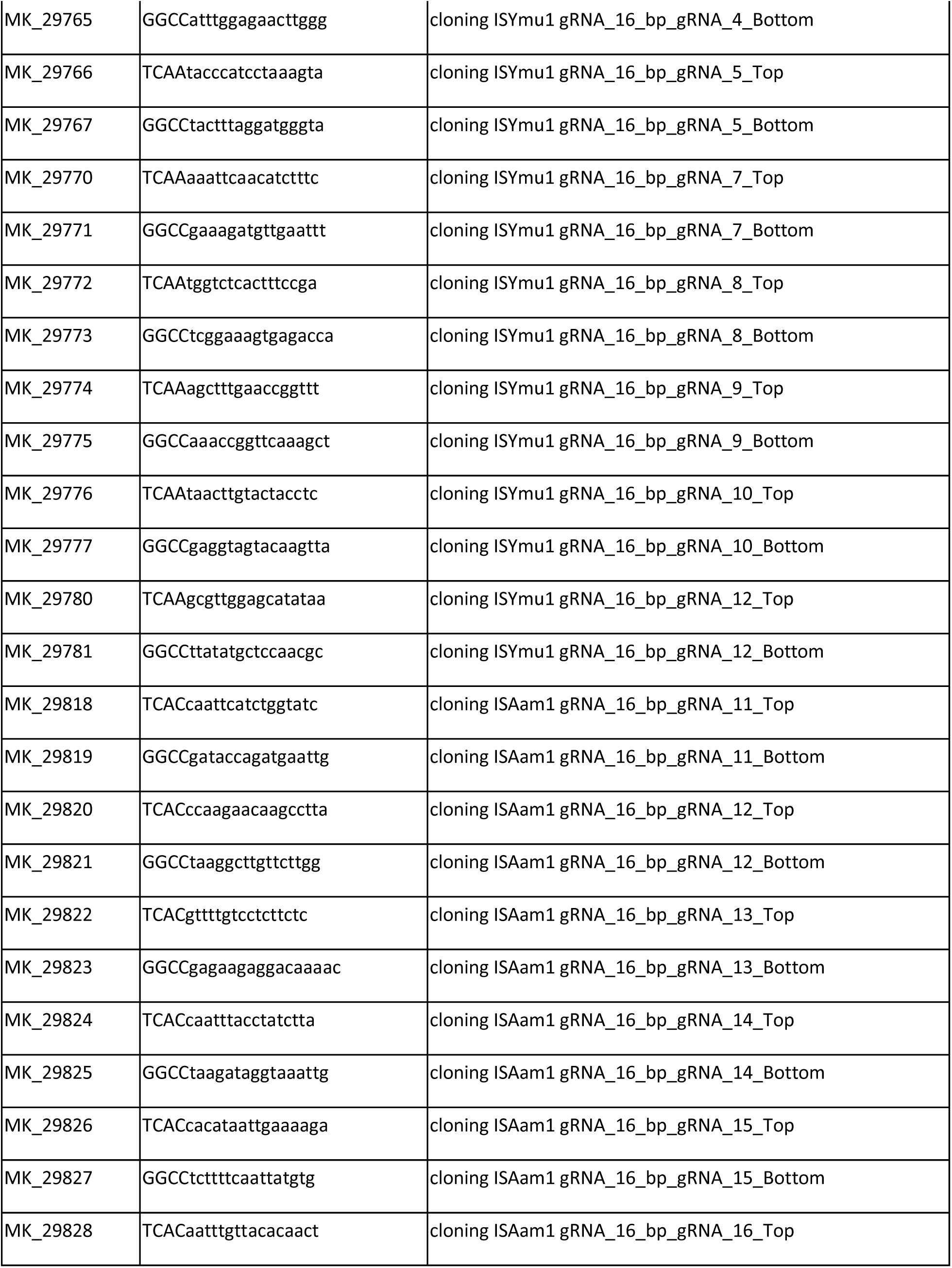
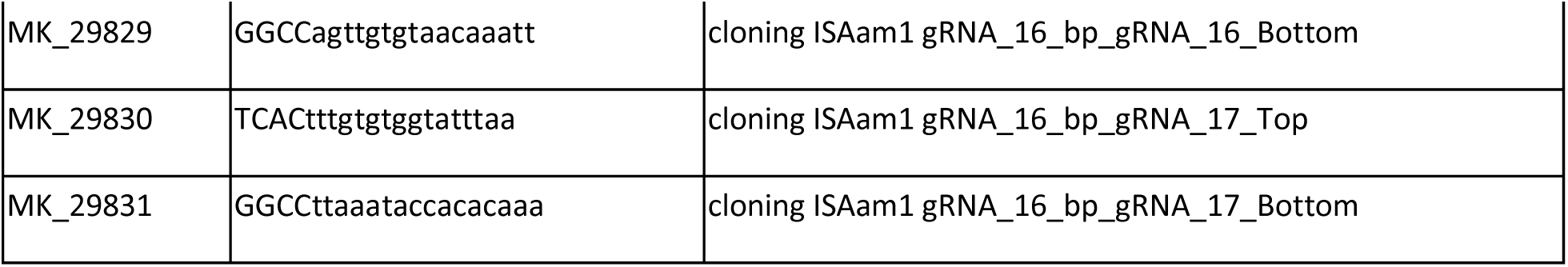
Oligos used for cloning gRNA sequences into intermediate TnpB plant expression vectors. The Oligo name column is an internal name for each oligo used. The Oligo sequence column lists the sequence of each oligo used for cloning the gRNA. The Oligo description column provides details for cloning each TnpB gRNA target site.

**Supplementary Table 8:**
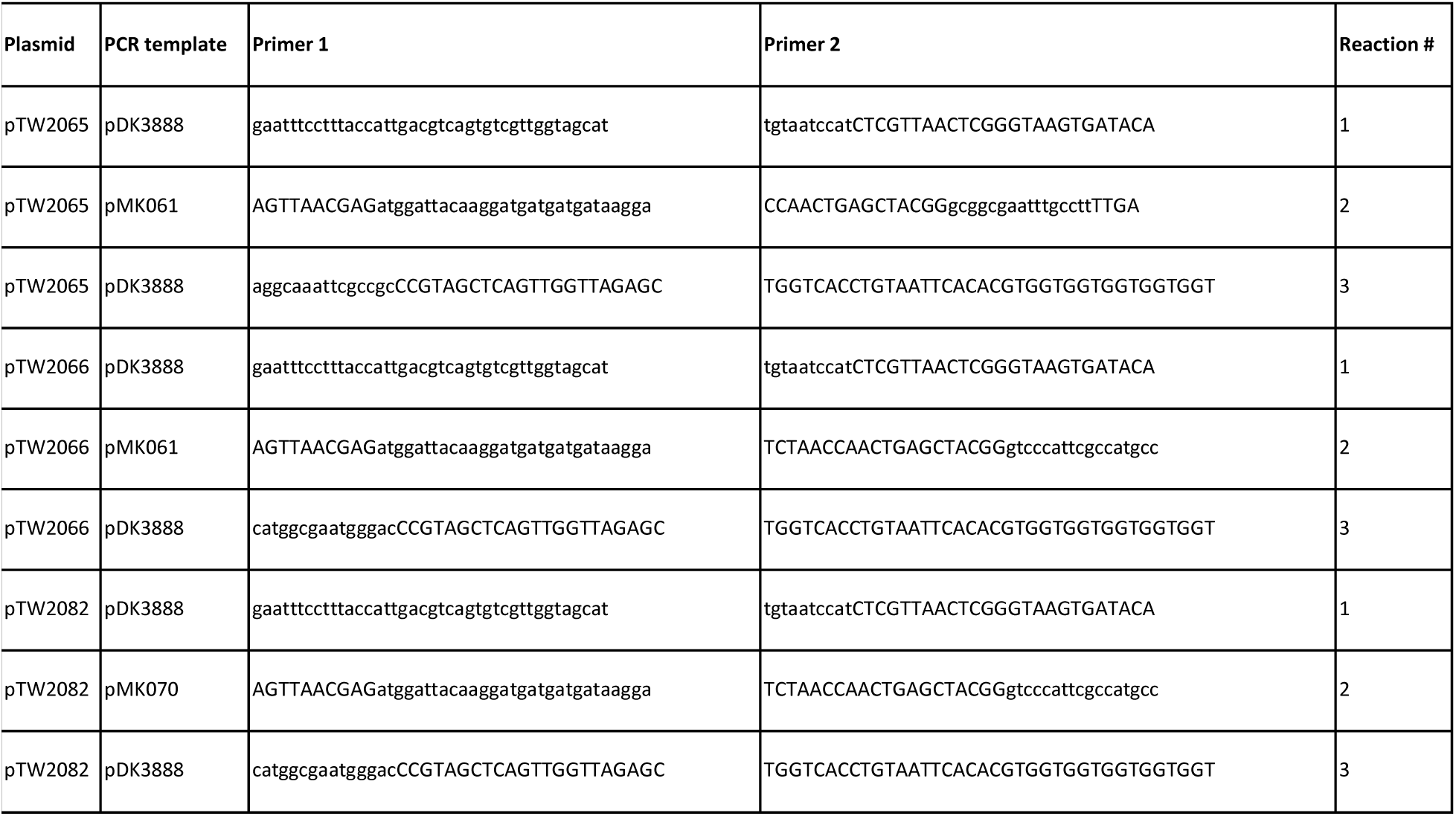
Cloning TRV Cargo Architecture_A and Architecture_B plasmids. The Plasmid column lists the name of each plasmid created. The PCR template column lists the name of each plasmid used for PCR DNA template. The Primer 1 and Primer 2 columns provide the sequence used to amplify fragments using the corresponding PCR template. The reaction column indicates the individual reaction for each PCR. The three PCR reactions, along with the restriction enzyme digested pDK3888, were used for NEB Hifi assembly to create the TRV2 vector listed in the Plasmid column.

**Supplementary Table 9:**
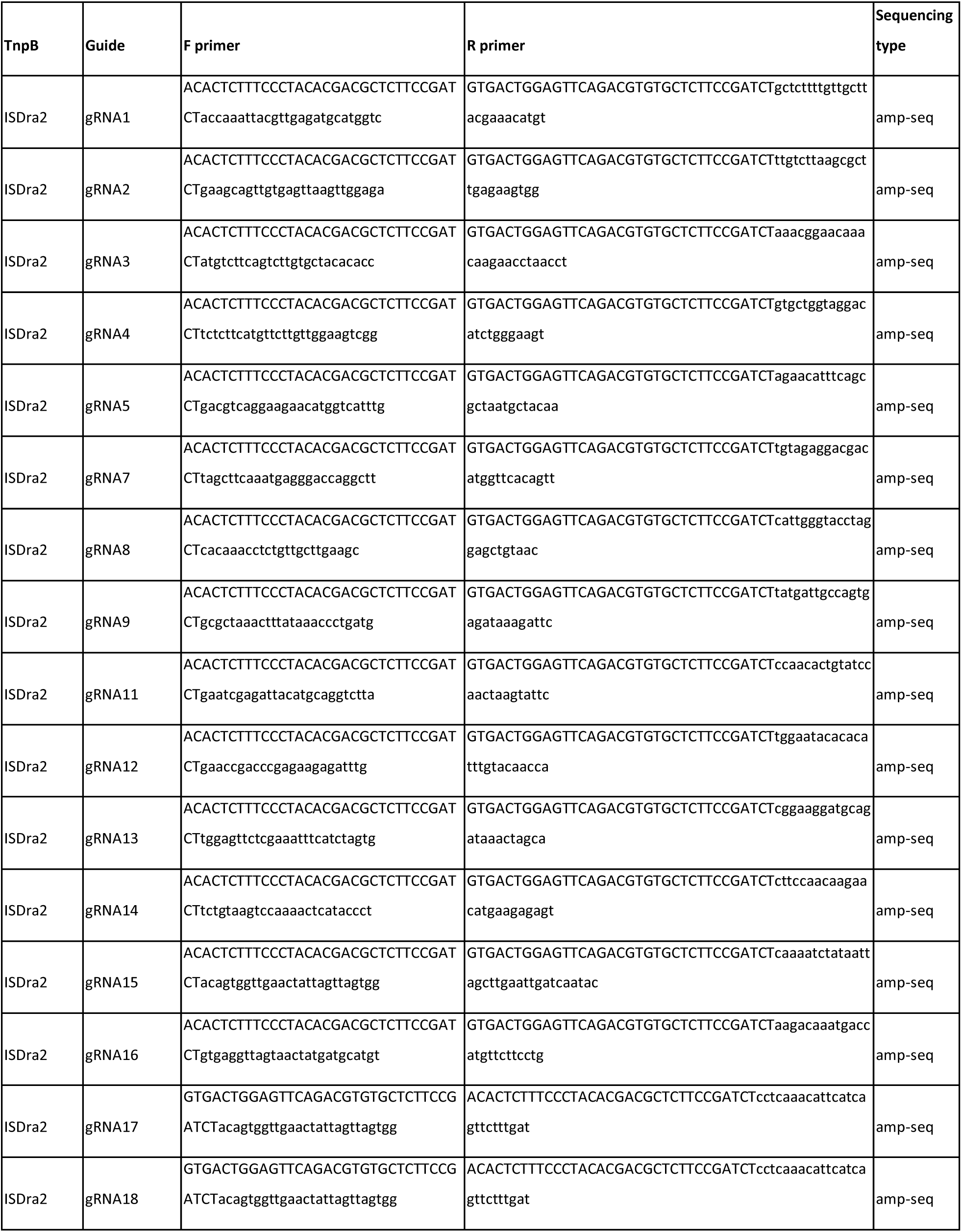
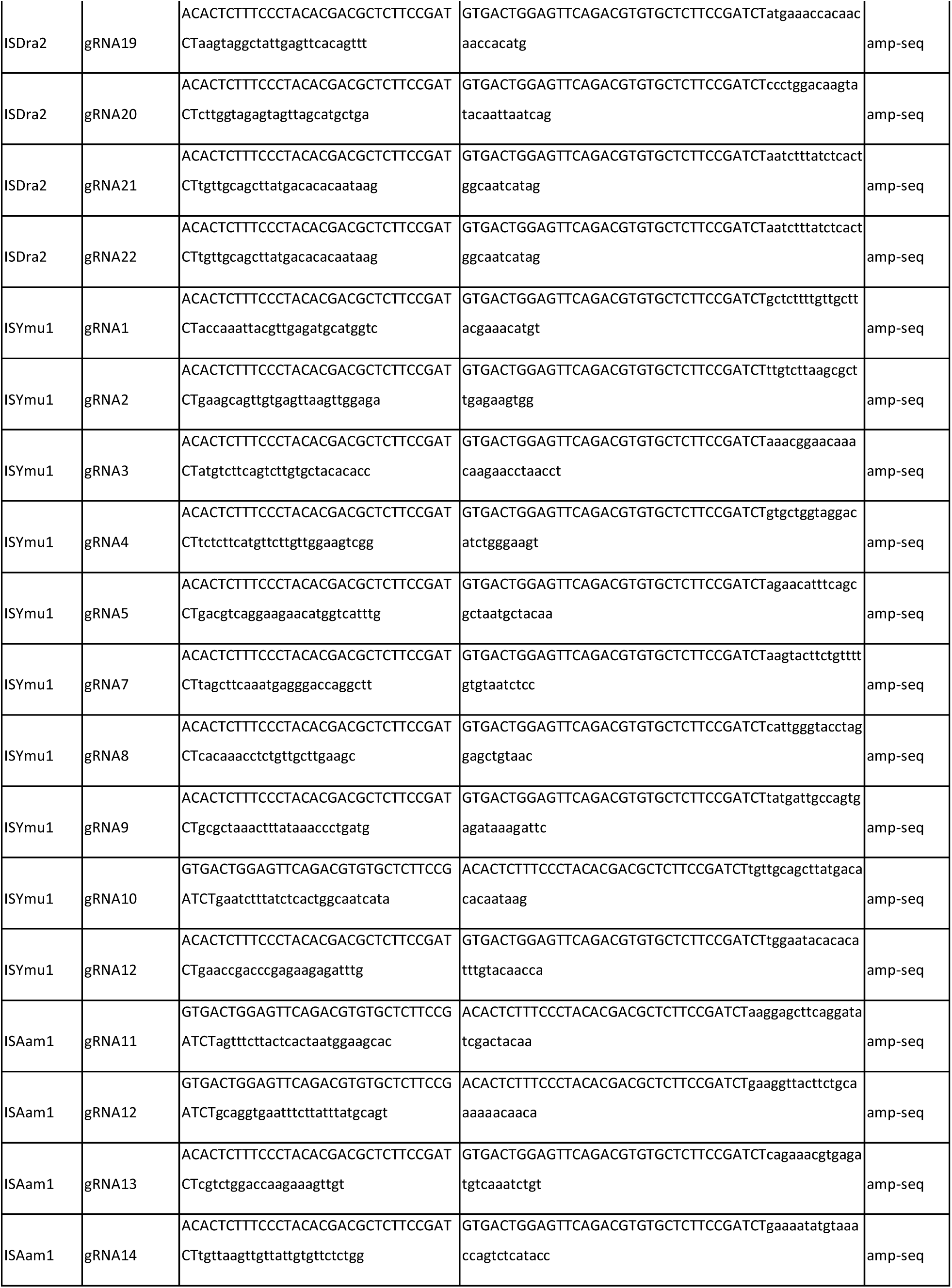
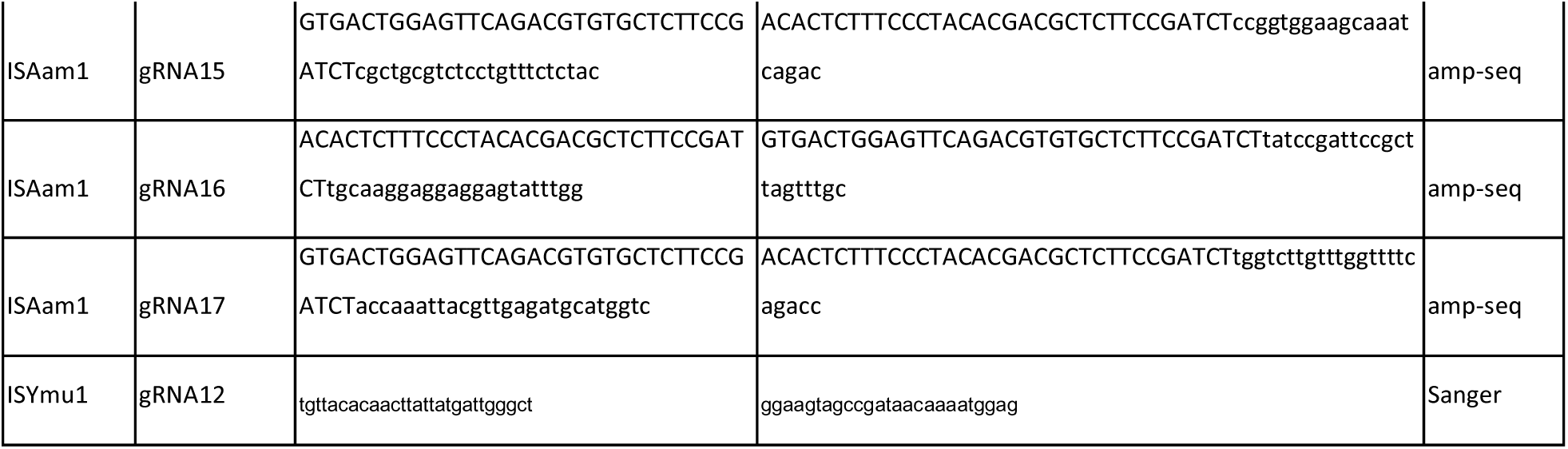
Primers used for amp-seq or Sanger sequencing. The TnpB and Guide columns indicate the site being targeted for each TnpB. The F primer and R primer columns list the oligo sequences used to amplify genomic DNA for amp-seq or Sanger Sequencing.

**Supplementary Table 10:**
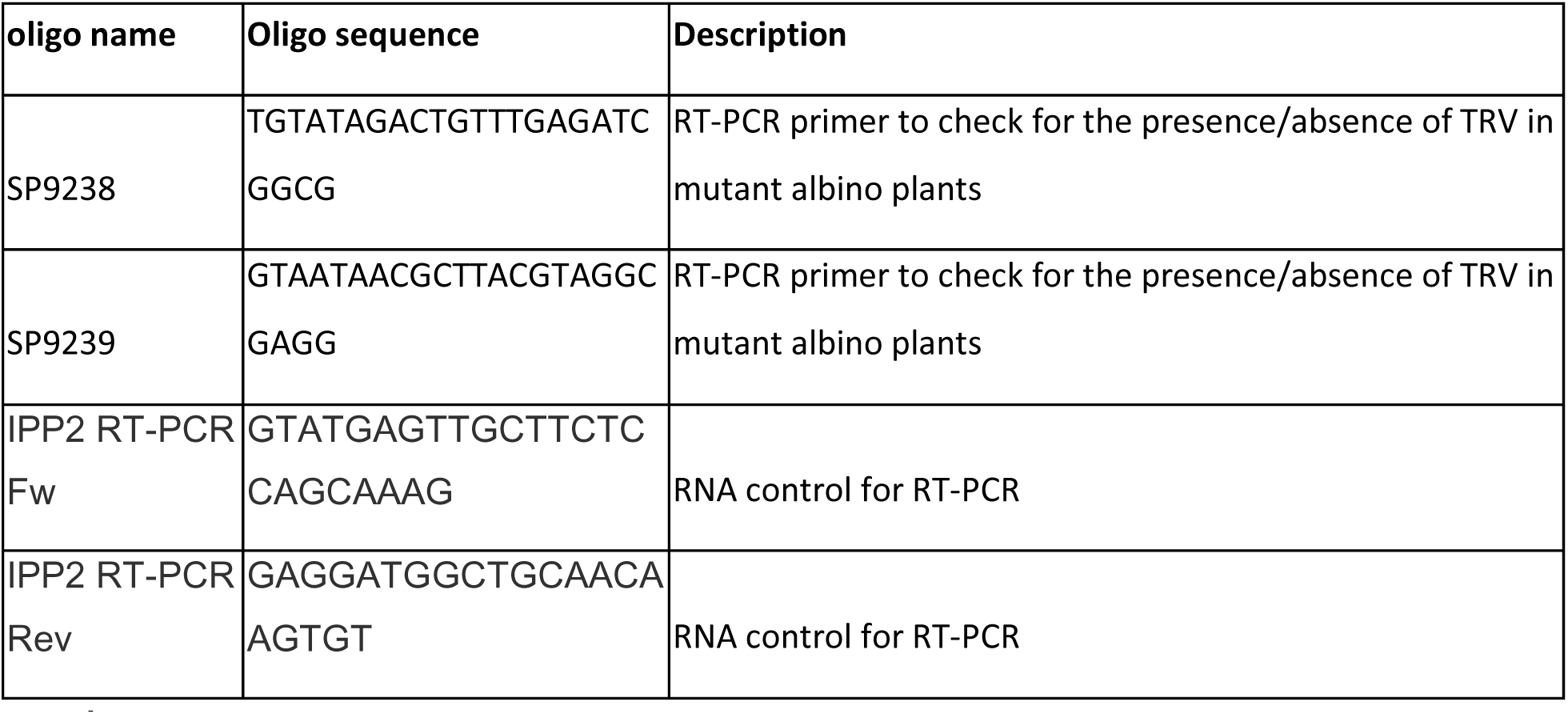
Oligos for RT-PCR. The name of each oligo is listed in the Oligo name column. Each oligo sequence is provided in the Oligo sequence column, with a description of what the oligos were used for in the Description column.

